# Humanized yeast to study the role of human ECE-1 isoforms in apoptosis in congenital heart disease

**DOI:** 10.1101/2023.06.19.545519

**Authors:** Hanhui Xie, Yan Huang, Edward J. Louis, Xiaodong Xie

## Abstract

Endothelin convert enzyme-1(ECE-1) plays a significant role in cardiovascular development including four isoforms with unclear function. Therefore, we are interested in studying the function of ECE-1 isoforms through mitochondria due to the high correlation between congenital heart disease (CHD) and apoptosis. Since the expression of human Bax and Bcl-xL in budding yeast (*Saccharomyces cerevisiae*) results in similar effects in mammalian cells, a yeast system was generated for mimicking human Bax-induced apoptosis and the expression of human ECE-1 isoforms was involved. The correlation between Bax-induced growth defect and the candidates of apoptosis via mitochondrial pathway was preliminarily investigated in this system. Furthermore, the phenotypes of ECE-1 isoforms have been identified through yeast growth defect. Individual ECE-1 isoform does not affect yeast growth but act as enhancers for the Bax-induced growth defect. ECE-1c is the strongest enhancer that affect the expression of candidates of outer membrane translocases. This study indicates that ECE-1 might play an important role in inducing apoptosis and we speculate these findings are possible to provide new perspectives with clarifying the mechanism of CHD.

## Introduction

ECE-1 is known as a group of significant regulators of vascular tone as a critical part of endothelin signal pathway. It is also reported that ECE-1 plays an important role in heart formation and the development of cardiac system (Robinson *et al*, 2014). Human ECE-1 is involved in proteolytic processing of precursors of endothelins (ETs). The precursors of ET are transformed to inactive big ETs by specific cleavage, and ECE-1 is the critical enzyme that regulate the step of converting ET-1 precursor to big ET-1 (Valdenaire *et al*, 1999).

The expression of ECE-1 through the period of adult development and was found in many different organs, particularly, it has a high level of expression in cardiovascular and endocrine systems (Korth *et al*, 1999). In addition, mutations of ECE-1 are also related to many diseases such as cardiac defects, Hirschrung disease and autonomic dysfunction (Hofstra *et al*, 1999). However, it is still unclear whether ECE-1 is correlated to apoptotic pathways. Currently, Researchers have found and defined four isoforms of human ECE-1, namely ECE-1a, ECE-1b, ECE-1c and ECE-1d. The existence and distribution of these four isoforms in cells has been identified and described. It is proved that ECE-1 isoforms are only have differences in their N-terminal region, which caused by a series of alternative splicing events. ECE-1a mainly expresses at the plasma and nuclear membrane, while ECE-1b only expresses intracellularly. The expression of ECE-1c is in another case that mainly occur on plasma membrane (Schweizer *et al*, 1997). The latest discovered isoform, which is ECE-1d, it widely distributes both cell surface and intracellular area. For example, it was detected at Golgi and several endosomal structures (Valdenaire *et al*., 1999).

Emerging evidence have shown that ECE-1 play a significant role in cardiovascular development and cardiac defects. An early study in 1998 has explored the role of ECE-1 during cardiac development with a murine model. In this study, ECE-1-null mouse embryos (ECE- 1−/−) was generated, which showed severe cardiovascular and craniofacial abnormalities and all died very soon with impaired breathing (Yanagisawa *et al*, 1998). Subsequently, Yanagisawa’s group continued this study with investigating the function of ECE-2 in cardiovascular development by generating ECE-2-null mouse model. The growth and development of ECE-2-null mice was normal and stay healthy in the whole lifespan. Interestingly, when the ECE-2-null mice were introduced into an ECE-1 knockout background by mating, the double knockout embryos showed a more severe cardiac defects on outflow structures than ECE-1 single knockout one, moreover, the double-null mice showed abnormal atrioventricular valve formation, which never be found in ECE-1-null embryos (Yanagisawa *et al*, 2000). One of the transcription factors named Nkx2.5, which play an important role during cardiac development was found to have a functional correlation with ECE-1a, ECE-1b and ECE-1c (Funke-Kaiser *et al*, 2003). In this study, H9c2 cells were transfected with an episomal expression vector for overexpressing Nkx2.5 and investigate the endogenous expression of ECE-1. The molecular level of ECE-1 was detected by using Dual-Luciferase Reporter Assay System and measured by relative luciferase activity (RLA) With the influence of overexpression of Nkx2.5, the RLA of all the three ECE-1 isoforms were upregulated. For further confirm the results and excluding the endogenous expression of Nkx2.5, the overexpression of Nkx2.5 was then conducted in endothelial EA.hy926 cells, the expression of three ECE-1 isoforms was also increased (Funke-Kaiser *et al*., 2003). Except for those studies of the association between ECE-1 and cardiac defects by using human and other mammalian cell line, epidemiological investigation was also widely used in studying the relationship of ECE-1 and CHD. For example, a case-control study with 945 CHD patients and 972 non-CHD controls was conducted by a Chinese group, who revealed two polymorphic sites were able to affect ECE-1 expression. These two variants were assessed by a combined analysis and found the appearance of 2-4 risk alleles (the ECE-1 338A and 839G) significantly increased the risk of occurrence of CHD, especially severe in females (Wang *et al*, 2012b).

Emerging evidence shows that apoptosis is an important event during cardiac morphogenesis including a large number of processes such as looping and formation of arteries, chambers (Poelmann & Gittenberger-de Groot, 2006). For example, Almost 80% of patients with CHD in their investigation were found to have a reduction of ACTC1 expression (Jiang *et al*, 2010). Interestingly, the inhibition of Bcl-2 expression is found in CHD patients who are associated with the abnormal expression of the *ACTC1* gene, and it is possible that the reduced expression of *ACTC1* could trigger the expression of caspase-3 and promote the apoptosis of cardiac tissues (Martin & Leder, 2001). In addition, there are also several mutations of *ACTC1* gene, which are correlated with cardiac defects from CHD patients but the functions of these genotypes remain unknown yet (Augiere *et al*, 2015). There is a general explanation for the correlation between genotypes of CHD and apoptosis. The abnormal phenotypes of transcription factors like GATA4 and structural proteins like ACTC1 caused by mutations and environmental factors, leading to inappropriate apoptosis might affects the development of cardiac tissues, finally leading to the onset of CHD (Kaichi, 2010). However, the mechanism between these genotypes and apoptosis is still unclear.

Budding yeast (*Saccharomyces cerevisiae*) has been widely used to study functions of human genes based on its clear genetic background and ease in handling and culturing. The life cycle of yeast is short with lower requirements for growth (Mager & Winderickx, 2005). Particularly, the most important advantage of yeast is its ability to put individual genes into yeast for expression makes it easier to investigate individual isoforms of human genes, which is not able to do in a human or other mammalian cell culture system due to a huge number of isoforms in human cells. Therefore, the methods of heterologous expression of human genes in yeast for screening possible disease-related factors or drug targets were emerging, even without considering any orthologs (Laurent *et al*, 2016).

Yeast apoptosis was first observed and described in the cells that contain a *CDC48* gene mutation. Many researchers thought yeast had no apoptosis prior to this. The *CDC48* gene has a homologue in mammalian cells (the *VCP* gene), the mutation of which can lead to apoptosis related behaviours (Madeo *et al*, 1997). Yeast apoptosis is normally caused by external factors including the stimuli of chemicals such as Acetic acid and the killer toxin that secret by double-stranded RNA viruses, while internal factors are include chronological aging and the heterologous expression of human apoptotic genes (Mazzoni & Falcone, 2008). Both the external and internal factors transmit cell death signals via the accumulation of reactive oxygen species (ROS). This is a critical pro-apoptotic process that plays an important role in the central regulation of yeast apoptosis. It relates to many pathways of PCD in yeast as a pro-apoptotic factor. Some representative inducers of human apoptosis such as caspase-8 and caspase-10 are not found in yeast, although there is a caspase-like protein called yeast caspase-1 (YCA1), which is a homologue to a human initiator caspase. YCA1 is the most important apoptotic inducer in yeast cells, it encodes the yeast caspase 1 protein (Yca1p), which functions as human caspase-8. Yca1p is usually be activated by external stimuli and induce apoptosis. For example, apoptotic stimuli from acetic acid, H_2_O_2_ and virally encoded killer toxins. Yca1p was activated by H_2_O_2_ stimuli through the generation of ROS, however, Yca1p plays a less important role during acetic acid induced apoptosis according to a previous study of flow cytometry, it is reported that only 20-30% caspase-positive cells under the acetic acid induced apoptosis (Saraiva *et al*, 2006). For the killer toxins, in the case of a low concentration, the apoptotic cell death is mediated by Yca1p and ROS accumulation. However, a high concentration of killer toxins results in necrosis, which is independent from Yca1p and ROS (Reiter *et al*, 2005). It is known that caspase 8 induces apoptosis by activating caspase-3 and caspase-7 in mammalian cells, whereas the cellular targets of Ycap1 in yeast remain unknown (Mazzoni & Falcone, 2008). Nuc1p is another important apoptotic factor in yeast, with human homologue of EndoG, and is located in the mitochondria. When yeast cells are under apoptotic stimulation, ROS accumulation induces the translocation of Nuc1p from the mitochondria to the nucleus (Büttner *et al*, 2007). Nuc1p functions by interacting with the H2B histone protein inside the nucleus to induce apoptosis.The human Bax inhibitor 1 (BI-1), which acts as a suppressor of apoptosis induced by Bax, can also have anti-apoptotic effect on Bax-mediated cell death in yeast (Madeo *et al*, 2004). The expression of human caspase-8 and caspase-3 are also carried out in yeast. It is reported that the expression of caspase-3 inhibits the growth of yeast cells, while the expression of caspase-8 results in yeast cell death (Kang *et al*, 1999). These findings are preliminary evidence that yeast may have a series of apoptotic processes.

The heterologous expression of mammalian Bcl-2 family in yeast has been investigated by several research groups for a long time. In 1995, one group transformed mouse Bax and Bcl-2 into yeast and conducted a viability assay by plated the transformed strain on galactose-containing media and glucose-containing media and to measure the number of colonies. It was reported that Bax is able to suppress the growth of yeast cells, while Bcl-2 makes the yeast cells survived from this suppression through interacting with Bax (Hanada *et al*, 1995).

Subsequently, another research that involved the heterologous expression of both murine Bax and Bcl-xL in yeast cells, the cell density was measured with OD_660_ at several time points, the data revealed that Bcl-xL inhibit the Bax-induced toxicity (Tao *et al*, 1997). However, these studies have no evidence to define the effect caused by mammalian Bcl-2 family proteins as apoptosis. Next year, another group conducted the heterologous expression of murine Bax and Bcl-xL in yeast and analysed the growth of cells by electron microscopy and tunnel assay, a series of morphological changes similar to apoptosis were observed with the overexpression of Bax in yeast cells, while the overexpression of Bcl-xL suppressed these changes (Ligr *et al*, 1998). In addition, mitochondrial activities like the increasing of cytochrome c (cyt-c) was found by the heterologous expression of human Bax (Manona *et al*, 1997). It indicates that the regulators of yeast apoptosis may have homologues in the human intrinsic pathway. The releasing of cyt-c is the main symbol of Bax-induced apoptosis in yeast. A channel of mitochondrial outer membrane named mitochondrial apoptosis-induced channel (MAC) has been found, which is affected by human Bax expression and get cyt-c through the membrane. Therefore, a ‘two steps’ model was generated from the discovery of MAC, this hypothesis demonstrated that Bax is able to insert into the mitochondrial outer membrane first during its expression in yeast, then the MAC is formed by inserted Bax that induce the release of cyt-c (Pavlov *et al*, 2001). It is also reported that yeast cells do show the morphological features of PCD, which are similar to human apoptosis such as the plasma membrane shrinkage, chromatin condensation, fragmentation and DNA breakdown (Madeo *et al*., 2004). Although these studies improve the understanding of the heterologous expression of mammalian Bcl-2 family, the mechanism is still unclear.

The emerge of this theory made yeast become a proper tool for studying the interaction between Bax and mitochondrial membrane proteins during apoptosis and let us pay attention to several mitochondrial genes that encode the translocases of mitochondrial membranes, though there is no evidence yet that the formation of MAC is related to mitochondrial translocases. The mitochondrial translocases of mitochondrial membranes, which including two types of complexes: The Translocases of Outer Membrane (TOM complexes) and the Translocases of Inner Membrane (TIM complexes). The TOM complexes are composed of Tom20 and Tom70, which act as two primary receptors. the imported proteins are recognized by these receptors and pass the outer membrane through the core of TOM complexes, which also known as the General Import Pore (GIP) to the IMS. Another five components of TOM complex are Tom22, a secondary receptor and the pore-forming protein Tom40, three small proteins of Tom5, Tom6, Tom7 (Herrmann & Neupert, 2000). The cleavable preproteins and most non-cleavable precursors are recognized by Tom20 and Tom70. Then the preproteins are delivered to the secondary receptor Tom22, which strongly bind to Tom40, the channel-forming protein that mediates the transport of preproteins. Tom5, Tom6 and Tom7 are relatively not necessary for the function of Tom complex, however, some researchers revealed that Tom5 is likely to assist the transportation of preproteins to Tom40. Tom6 plays a role in maintaining the stability of Tom complex (Dietmeier *et al*, 1997). Tom7 may have an opposite function compared with Tom6, in order to allow the structure of the complex be flexible, a changeable conformation for better performing its function of protein transport (Wiedemann *et al*, 2004). In order to transport the proteins through the outer membrane, it is also essential to rely on β-barrel protein, which is a kind of precursor protein assembled by the sorting and assembly machinery (SAM) complex. The β-barrel protein is a kind of small signalling protein for assisting protein transport, membrane anchors and defending against aggressive proteins (Schulz, 2000). It also helps to maintain the integrity of the outer membrane and pore forming (Wiedemann & Pfanner, 2017). SAM complex includes 4 subunits: Sam50, Sam35, Sam37 and mdm10. Sam50 is the main component of SAM complex and mediate the formation of β-barrel protein. Sam35 and Sam37 are the secondary outer membrane proteins for maintaining the function of Sam50 (Laan *et al*, 2006). As the translocases of inner membrane, TIM complex drives its function with TOM complex for sorting the proteins from outer membrane and translocate proteins to the matrix. The most well-studied component of TIM complex is Tim23, which is the main channel-forming component. Tim17 is also necessary as another important component of the membrane-integrated channel, which contributes to assist Tim23 to fulfil the function of importing proteins to the matrix (Laan *et al*., 2006). Tim50 and Tim21 are two subunits of Tim23, which can be crosslinked with the precursor proteins from TOM complex and transfer them to Tim23. Tim21 is the latest identified component that interact with Tom22 and deliver the signal of protein transport and sorting to the channel of Tim23 (Chacinska *et al*, 2005).

Based on the studies about the possibility of CHD, the importance of ECE-1 in cardiac development and the description of mitochondrial translocases above, we hypothesize that pathogenesis of CHD might be associated with apoptosis, and ECE-1 isoforms act as potential regulators of apoptosis during the development of CHD. However, the apoptotic mechanisms among four ECE-1 isoforms are understood incompletely through these studies. Moreover, the understanding of the function of ECE-1 during the development of CHD is limited. Therefore, we are interested in studying whether the four isoforms have different function, especially in inducing apoptosis through mitochondrial pathway. Depend on these findings, we hope to understand the possible function of ECE-1 isoforms in CHD.

As yeast is a powerful tool for studying human apoptosis, we generate a human Bax/Bcl-xL expression system with budding yeast for mimicking the apoptotic effect and confirm the phenotypes by spot assay, then to investigate the mechanism of Bax-induced growth defect by measuring the mRNA expression level of several apoptotic factors and mitochondrial genes in yeast, which including *YCA1, NUC1, BI-1, Tom40, Tom20, Tom70, Tom22, Tom5, Tom6, Tim50, Tim17, Tim21, Tim23, Sam35, Sam37* and *Sam50*.

Four ECE-1 isoforms are transformed into Bax/Bcl-xL expression system subsequently for investigating the roles of individual isoforms during Bax-induced growth defect. Firstly, to generate the strains contain both Bax and ECE-1 isoforms for confirming the phenotypes of ECE-1 isoforms during Bax-induced growth defect. Next, to explore whether individual ECE- 1 isoforms have effect on the mRNA expression level of candidate genes we listed above. Finally, the mRNA level of these mitochondrial genes needs to be measured under the expression of both Bax and ECE-1 isoforms.

## Results

### The new yeast system for mimicking apoptotic effect is successfully generated

The haploid strain contains contains Bax (W303a: pRS405/Bax, which was presented as ‘Bax’ in the further figures) and empty vector of Bax (W303a: pRS405/Gal1p, which was presented as ‘Vehicle’ in the further figures) was mated with W303 α: pRS424/ Bcl-xL and W303 α: pRS404/ Bcl-xL (presented as ‘Bcl-xL’ in the further figures) respectively, and also mated with W303α: pRS424/Gal1p-MS and W303α: pRS404/Gal1p-MS (The negative controls, which were presented as ‘NC’ in the following figures) respectively. Next, the colony PCR was conducted with eight diploid strains generated from yeast mating for confirming the existence of Bax, Bcl-xL and the mating type, three positive clones of each strain were picked and used as templates. Therefore, 24 PCR reactions of each gene were shown (Figure 1). All the bands of PCR products were presented in the correct size, that means the mating was successful and both Bax and Bcl-xL were involved in the new diploid strains. Then a spot test was conducted with the LEU-TRP drop-out agar media with adding both glucose (Glu+) and galactose (Gal+) for switching off/on the expression of Bax/Bcl-xL for mimicking the apoptotic effect of Bax and anti-apoptotic effect of Bcl-xL in yeast (Figure 2). The results indicated that the new constructed Bax/Bcl-xL yeast system works correctly, the Bax-induced growth defect in yeast was presented as expected, the anti-apoptotic effect of Bcl-xL was also observed when the Bcl-xL was integrated into pRS404. However, no significant alternation caused by Bax and Bcl-xL was shown in the pRS424 group.

**Figure 1.**
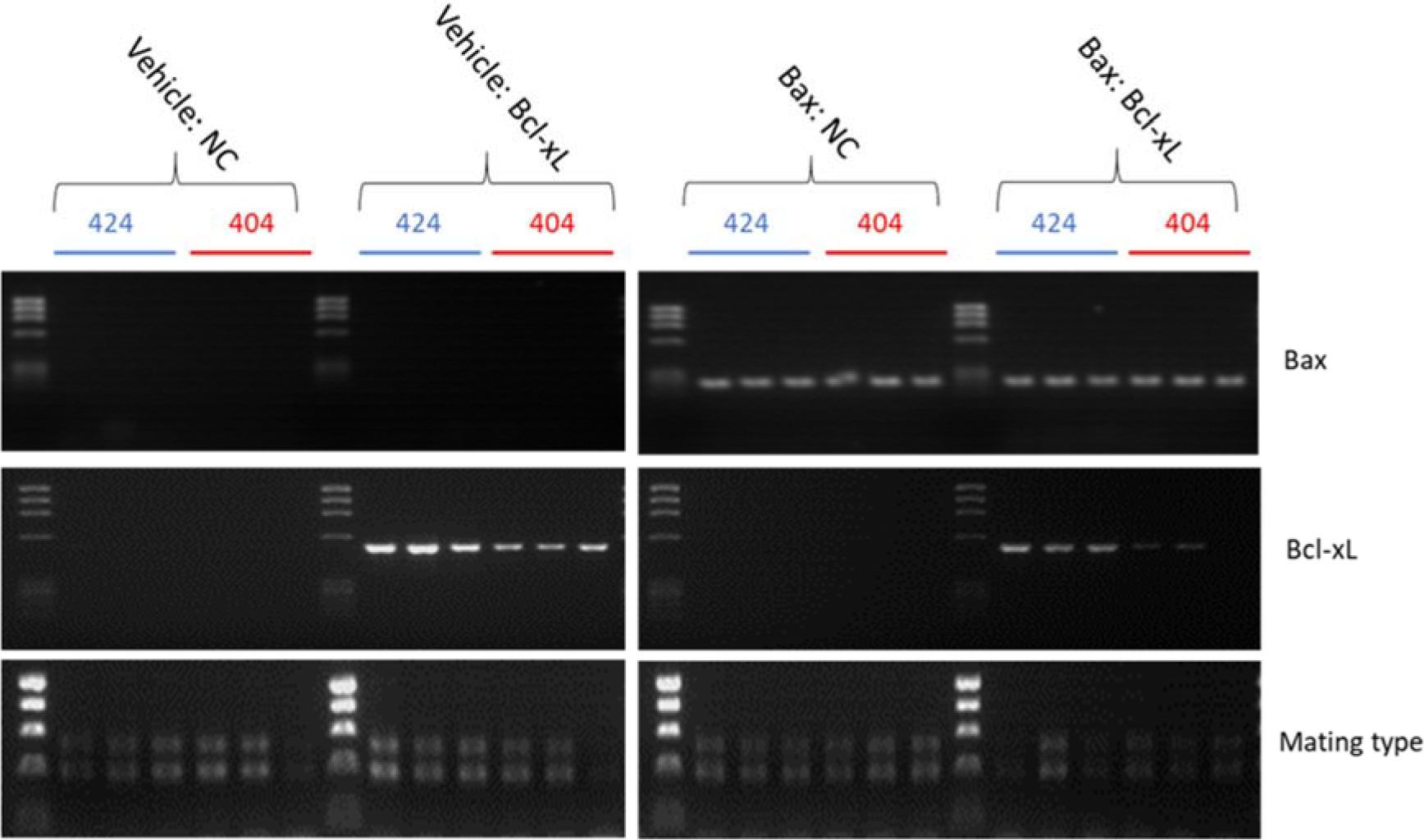
The identification of genotypes of diploid strains by colony PCR. This image indicates both Bax and Bcl-xL were successfully introduced into yeast cells. The appearance of both bands of MATa and MATα illustrates the diploid cells we need were successfully generated. The PCR products were separated by 1% w/v agarose gel.

**Figure 2.**
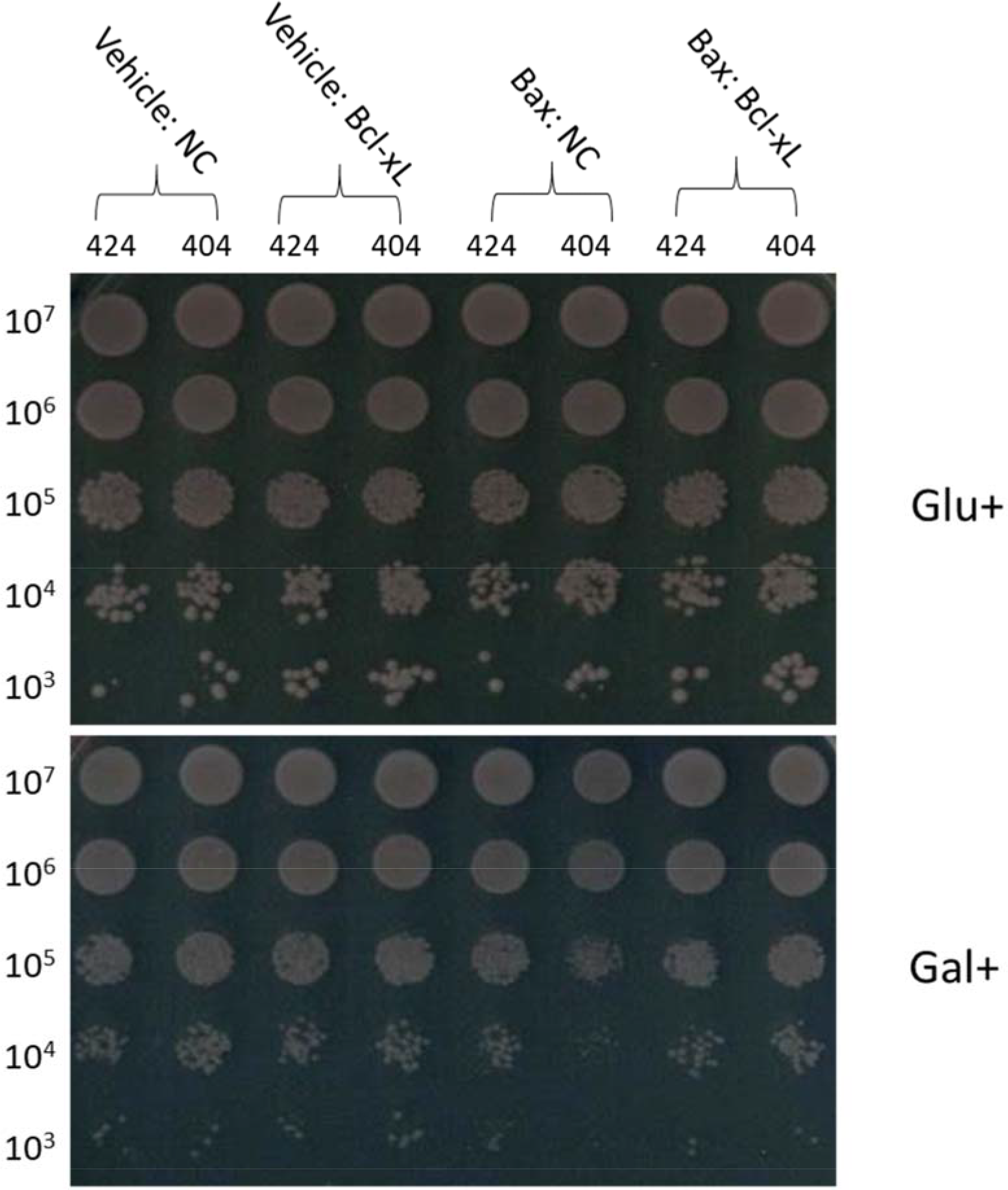
The phenotypes of diploids that contain both Bax and Bcl-xL. The plates were incubated at 30°C for 48h, the harvested cells were processed with 10-fold dilution, which ranging from 10^7^ cells to 10^3^ cells. The image showed when switching on the expression of Bax/Bcl-xL with galactose, a growth defect caused by Bax could be observed in yeast, this effect is able to be inhibited by Bcl-xL successfully with its integration into pRS404. While there was no obvious alternation in the strains that contain pRS424 when switching off/on the expression of Bax/Bcl-xL.

### The semi-qPCR analysis is available to investigate the mechanism of human Bax-induced growth defect in *S.cerevisiae*

The humanization of apoptosis in yeast was proved to be successful by doing colony PCR and spot test. Therefore, it is necessary to explore how could human Bax influence the growth of yeast cells or cause apoptosis in molecular levels. In fact, Bax-induced cell death in yeast is involved in mitochondrial system have been approved and demonstrated by many researchers in the past few years. Following the heterologous expression of Bcl-2 family proteins was accomplished in yeast, it is demonstrated that the expression of a modified Bax, which contains a c-myc tag is able to cause much stonger release of cyt-c from yeast mitochondria (Manon *et al*, 1997). Besides, another three proteins apoptosis-inducing factor (AIF), smac/diablo and endonuclease G, which are all plays essencial role during apoptosis (Priault *et al*, 2003).The Endonuclease G is located in mitochondria, it has homologue in yeast named Nuc1p, which is an important regulator of yeast apoptosis by mediating DNA degradation directly (Burhans & Weinberger, 2007). Therefore, we suppose that yeast Nuc1p is involved in Bax-induced cell death. Moreover, there is also a lack of evidence that other mitochondrial membrane proteins like Tom complexes of outer membrane and Tim complexes of inner membrane were associated with Bax-induced growth defect. Based on these, to investigate the possible function of several representative mitochondrial genes during Bax-induced growth defect, or whether these mitochondrial genes were involved in the mechanism of Bax-induced growth defect will be the second step of our study.

The function of all the candidate genes will be assessed by the dectection of mRNA expression levels. Although the protein levels have been demonstrated to be more conserved and imformative than the mRNA levels for conducting gene expression studies, It has been identified that the assessment of mRNA levels are still able to be a significant proxy for explaining protein levels primarily (Laurent *et al*, 2010; Liu *et al*, 2016). The mRNA-protein relationship in yeast has been investigated by a large-scale proteome and transcriptome analysis, the comparison between mRNA and protein abundance indicated that proteins are difficult to be detected with low mRNA levels. In contrast, higher mRNA levels raise the detectability of proteins, the probability of protein abundance do not increase any further when mRNA abundance exceeds a specific value (Ramakrishnan *et al*, 2009; Vogel & Marcotte, 2012). Therefore, the detection of mRNA levels would be much efficient as a primary screening method when faced with a large numbers of candidate genes, the positive results would be more significant for conducting the further analysis of protein levels, while the negative results of mRNA levels would be less important for the further studies.

Based on the results of the new strains confirmation, the strains contain pRS404/Gal1p-MS and pRS404/Gal1p-Bcl-xL-MS were much obviously influenced by the heterologous expression of Bax/Bcl-xL. Therefore, four strains were chosen for doing the semi-qPCR, which were Vehicle: NC (404), Vehicle: Bcl-xL (404), Bax: NC (404) and Bax: Bcl-xL (404). The empty vector of pRS426 (will be used as the vector of ECE-1 isoforms in another group of results) was transformed into these four strains. These newly generated strains were named as: Vehicle:Vector::NC, Vehicle:Vector::Bcl-xL, Bax:Vector::NC and Bax:Vector::Bcl-xL (‘Vehicle’ represents pRS405, ‘Vector’ represents pRS426, ‘NC’ represents pRS404, they are all act as a control without any heterologous genes), which were screened by SD-LEU-URA-TRP agar media, then the cells of each strain were cultured with SG-LEU-URA-TRP liquid media for switching on the heterologous expression of Bax/Bcl-xL for 24h. The cells were finally harvested by centrifuging, then RNA and DNA extraction was conducted, the samples were stored at −80°C for further experiments.

For making sure the grey value can be used for measuring the concentration of PCR products, the plasmid DNA of pRS404/Bcl-xL is needed for generating a standard curve equation due to the good stability of plasmid DNA. First, PCR reactions was conducted by using pRS404/Bcl-xL as template with several dilutions, the concentration of PCR products was detected by nanophotometer and the grey value of each PCR products was measured by Image J. After that, the standard curve was made and the linear regression analysis was conducted by Graphpad Prism 7. The results showed that the curve fits the linear regression according to the value of R^2^ (Figure 4). *ACT1* is a commonly used housekeeping gene for budding yeast studies, which is act as the reference gene due to several reasons: The sequence of *ACT1* is highly conservative, which encode the structural protein including the components of cytoskeleton. It is difficult to alter the expression level of *ACT1* by most of external and internal factors (Cankorur-Cetinkaya *et al*, 2012; Teste *et al*, 2009). For confirming whether the mRNA level of *ACT1* is able to be changed under the heterologous expression of Bax/Bcl-xL, the cDNA samples of Vehicle:Vector::NC, Vehicle:Vector::Bcl-xL, Bax:Vector::NC and Bax:Vector::Bcl-xL were obtained from Reverse-transcription PCR, 20ng cDNA template was used for each PCR reaction. Here we assume that the efficiency of all the PCR reactions is same, therefore, the grey value of four bands of *ACT1* was measured, then the grey value of Vehicle:Vector::NC was set as 100% (divided by itself) for acting as a control, the grey value of another three was divided by the control respectively. The final data that express as the percentage represents the expression level of *ACT1* (Figure 3). The result indicates that the expression of Bax and Bcl-xL did not alter the expression level of *ACT1*. This means *ACT1* is a proper choice as a reference gene for the further semi-qPCR analysis.

**Figure 3.**
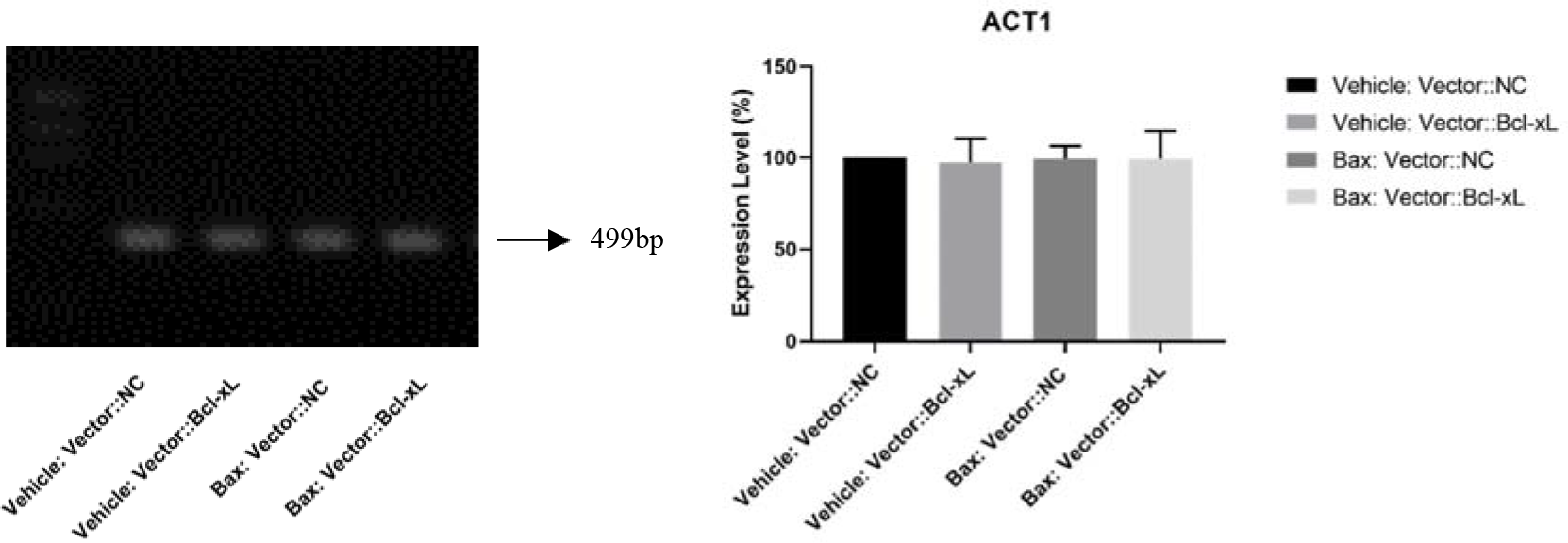
The expression level of *ACT1* under the heterologous expression of Bax/Bcl-xL. The image showed that the expression level of *ACT1* is remain to be same in all 4 different strains. The expression of *ACT1* was not influenced by either Bax or Bcl-xL. The experiment was repeated for three times and the data was presented as mean±SD. The grey value was measured by Image J. One-way ANOVA analysis was conducted by GraphPad Prism 7.

**Figure 4.**
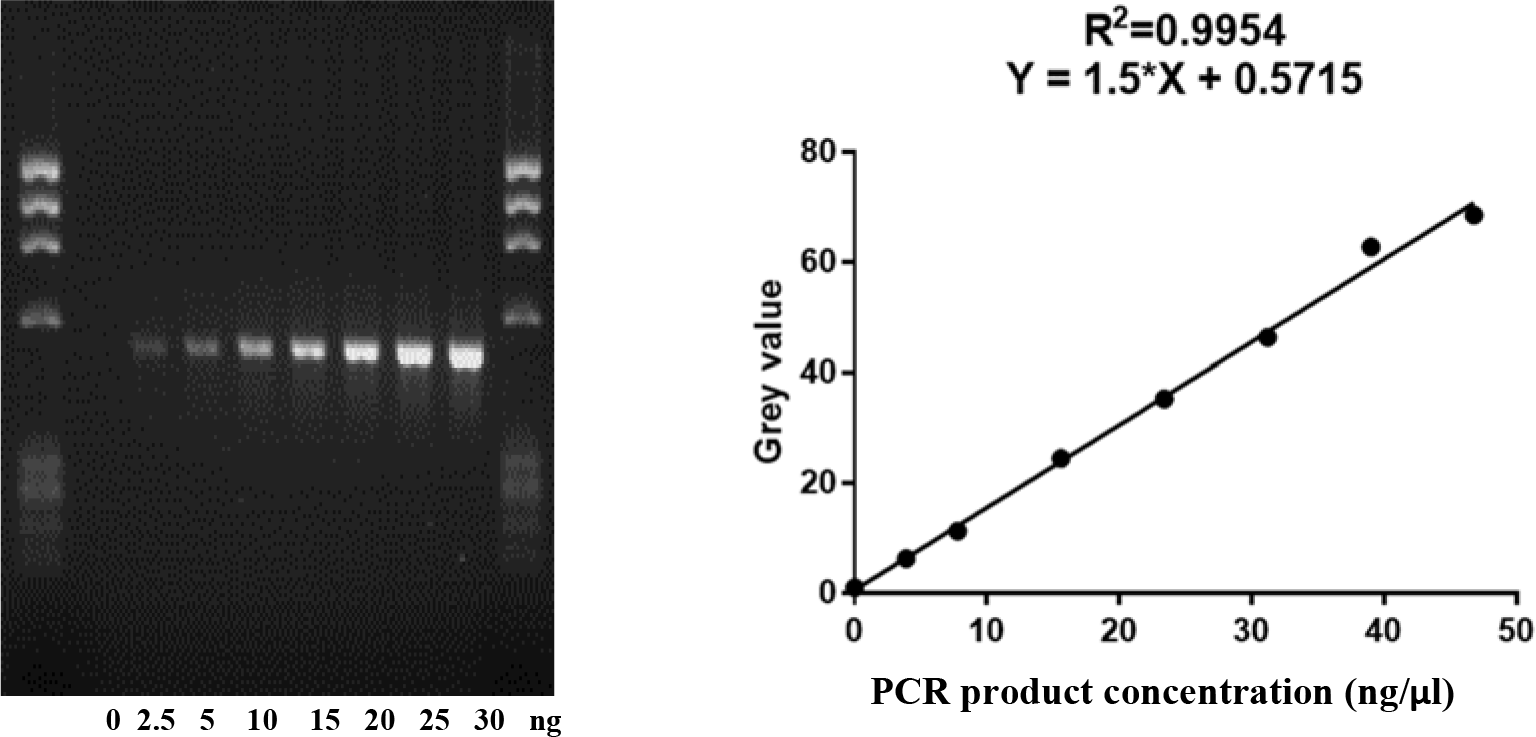
The linear regression analysis of PCR products of pRS404/Bcl-xL. The plasmid DNA was diluted to a concentration of 10ng/µl for using as the template with 2.5, 5, 10, 15, 20, 25, 30ng respectively. Bcl-xL 113Fw and Bcl-xL 632Rev were used as primers (shown in Table 1). 8 PCR reactions (a negative control PCR was included) were done. The PCR products were isolated with 2% agarose gel. The R^2^ of standard curve is approximately equal to 1, therefore the grey value is able to be used as a standard to estimate the concentration of PCR product.

**Table 1.**
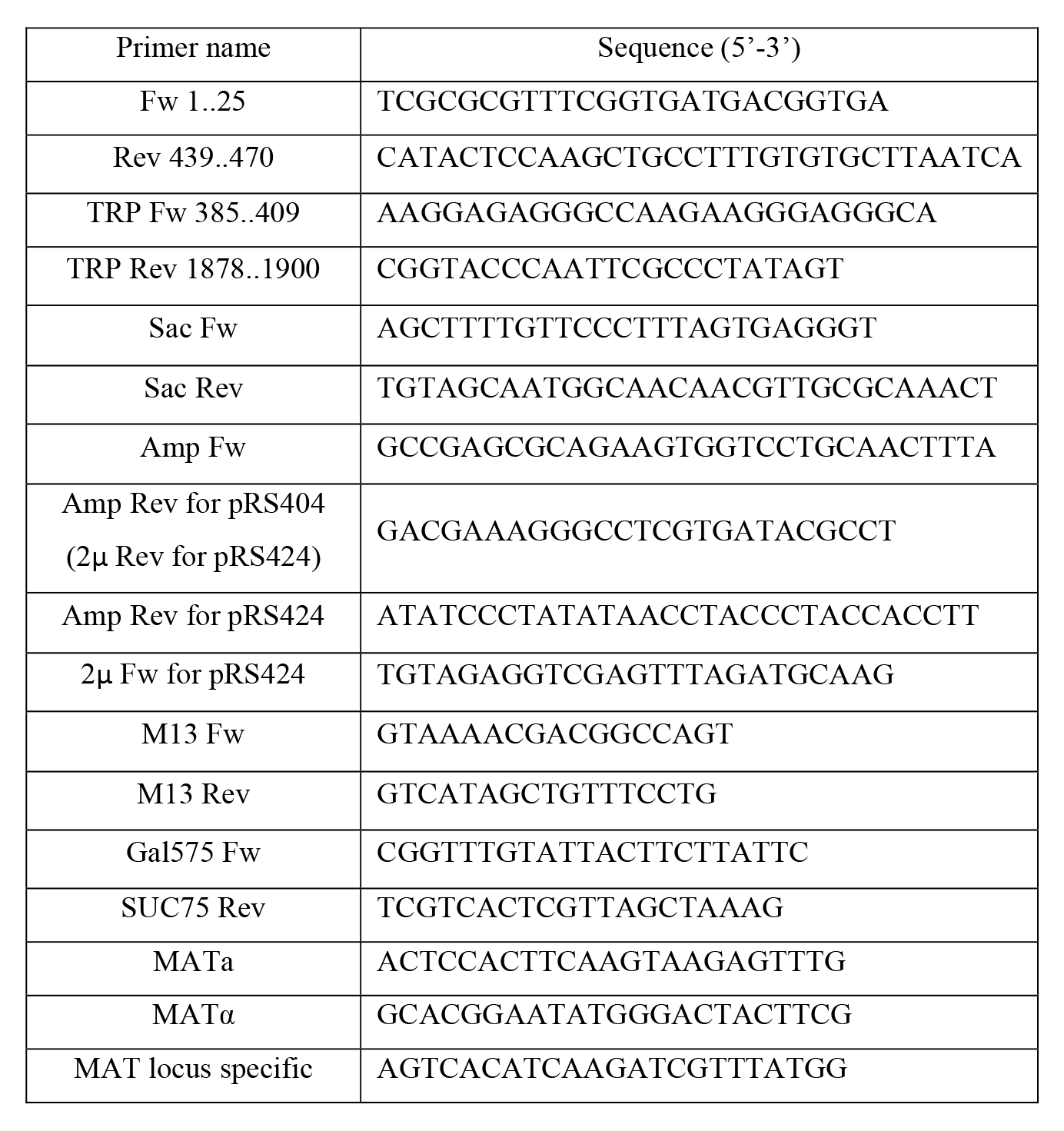
Primers used for construction and PCR confirmation

### Human Bax-induced growth defect decreased the expression level of YCA1 and NUC1

For explore whether the growth defect caused by human Bax in yeast is depend on canonical apoptotic pathway, we detected the expression level of YCA1, NUC1 and BI-1 with switching on the expression of human Bax and Bcl-xL. cDNA and DNA samples were diluted to 10ng/ µl and use 20ng for each PCR reaction as the template. PCR reactions of DNA performed as a control group. The results showed that the expression of YCA1 is not activated but inhibited by the individual expression of human Bax and Bcl-xL, when the expression of both Bax and Bcl-xL was triggered, it increased slightly. The expression level of another pro-apoptotic regulator NUC1 is also decreased by the individual expression of Bax and Bcl-xL, however, it recovered almost to the normal level under the expression of both Bax and Bcl-xL. As a representative anti-apoptotic molecule, the expression level of yeast Bax inhibitor-1 was almost not change under the expression of human Bax and Bcl-xL (Figure 5).

**Figure 5.**
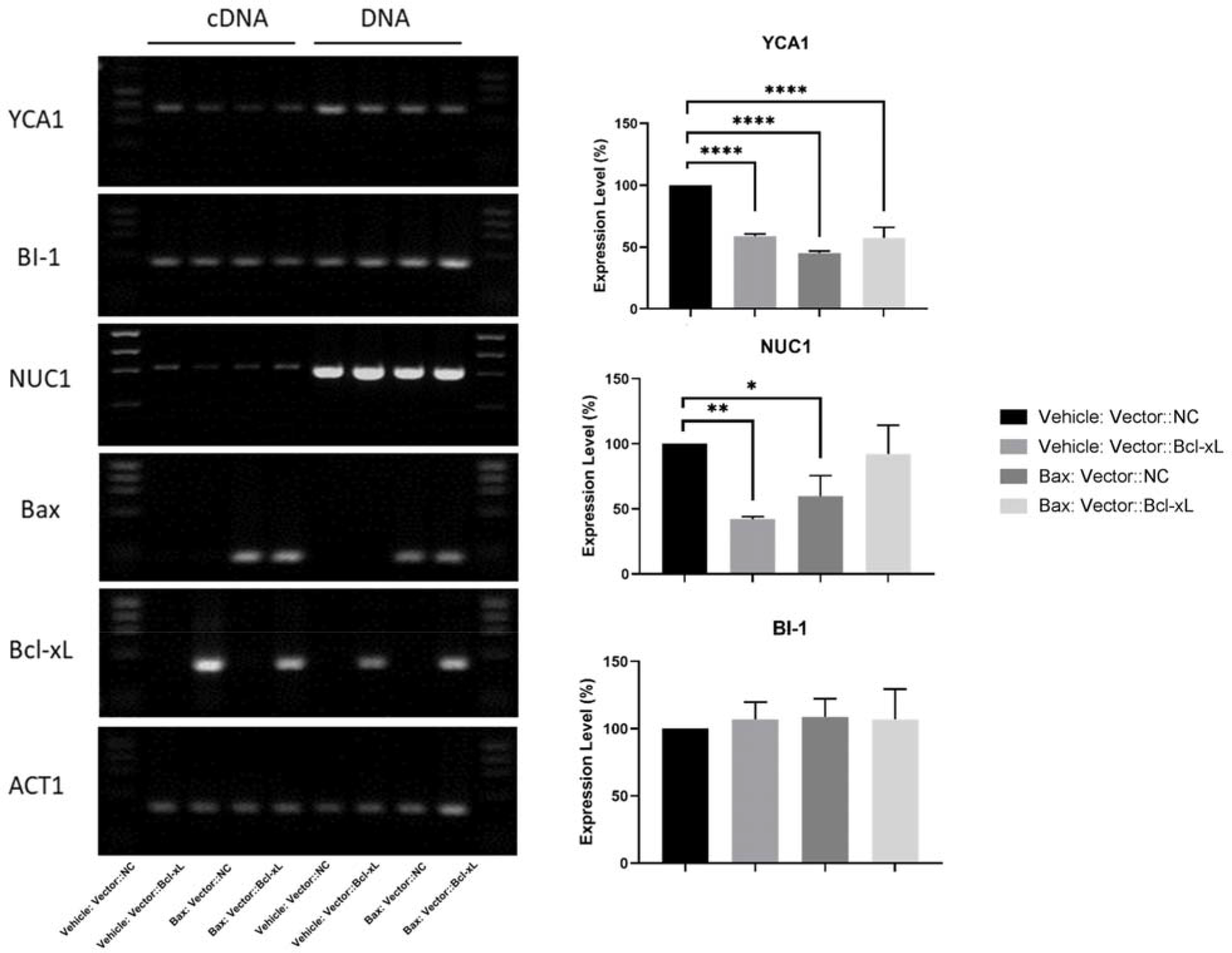
The expression level of three representative yeast apoptotic molecules under the heterologous expression of Bax/Bcl-xL. The images showed that YCA1 expression is decreased by the expression of both human Bax and Bcl-xL. However, the expression of NUC1 is inhibited only by Bcl-xL. The expression of yeast BI-1 was not altered by human Bax and Bcl-xL. The PCR products were isolated by 1.5% agarose gel. Gene expression level was measured by the ratios of the grey value of the bands of detected genes and reference gene. The experiment was repeated for three times and the data was presented as mean±SD. Grey value was measured by Image J and One-way ANOVA analysis was done by GraphPad Prism 7 (*p<0.05, **p<0.01, ***p<0.001, ****p<0.0001).

### The expression of human Bax and Bcl-xL have no effect on TOM complex

TOM complex plays an important role in general importing and sorting proteins to the inner membrane and matrix. It has been reported that Tom22 act as a receptor during the Bax-dependent apoptosis (Bellot *et al*, 2007), however, there is no clues about other components of TOM complex that related to Bax-induced growth defect. Therefore, we hypothesize that human Bax-induced growth defect is involved in the protein importing pathway mediated by TOM complex. The expression of 6 components of TOM complex were detected with switching on the expression of human Bax and Bcl-xL. Unfortunately, none of these members of TOM complex was significantly influenced by the expression of Bax/Bcl-xL. The expression of Tom70 is slightly decreased by the expression of human Bax. In contrast, the expression of Tom22 seems increased slightly with the expression of both Bax and Bcl-xL. The expression of Tom40, Tom20 and another two small Toms of Tom5 and Tom6 has no significant change (Figure 6).

**Figure 6.**
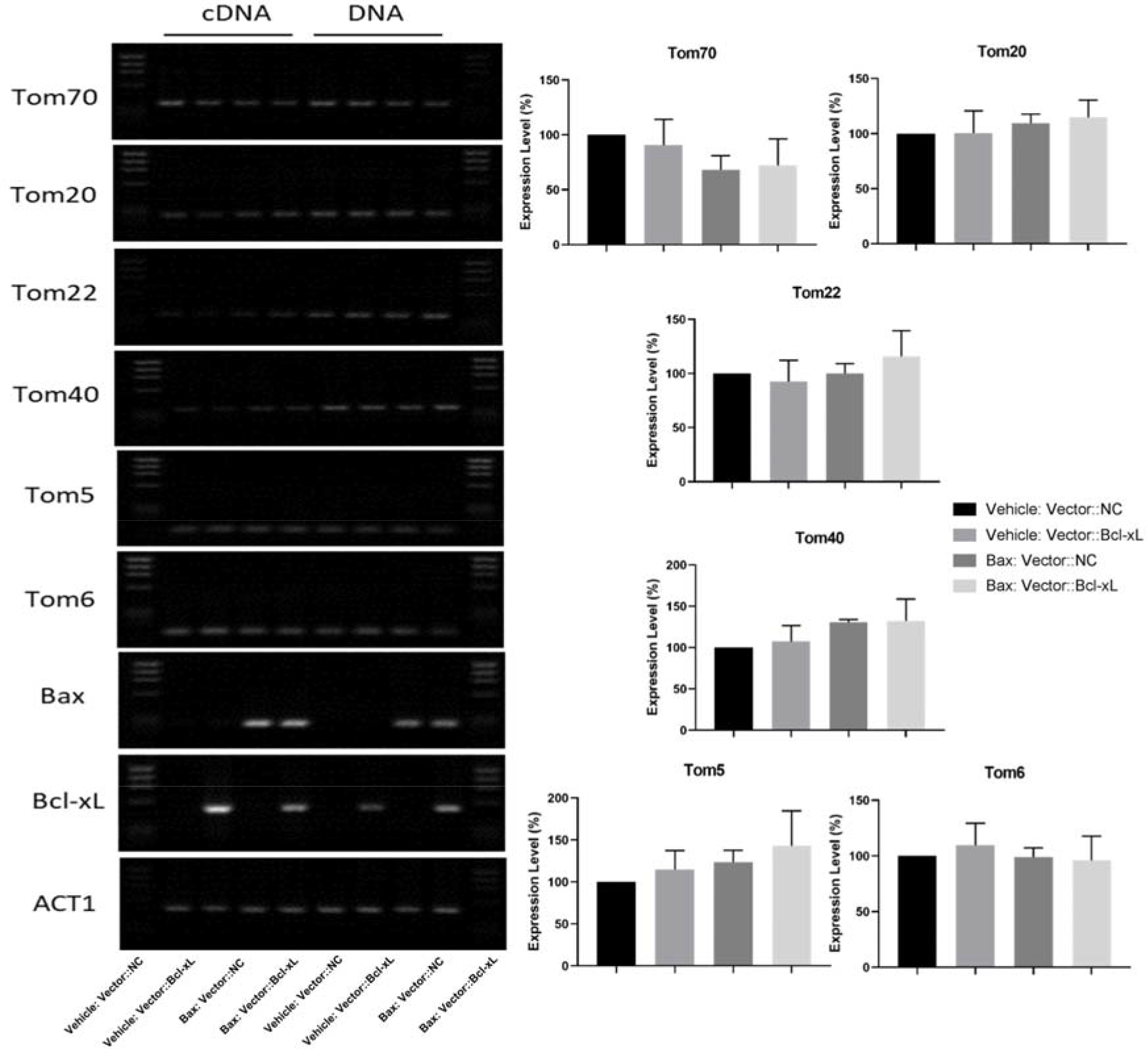
The expression level of representative subunits of TOM complex under the heterologous expression of Bax/Bcl-xL. The images showed that human Bax/Bcl-xL did not alter the expression level of any gene of TOM complex. The PCR products were isolated by 1.5% agarose gel. Gene expression level was measured by the ratios of the grey value of the bands of detected genes and reference gene. The experiment was repeated for three times and the data was presented as mean±SD. Grey value was measured by Image J and One-way ANOVA analysis was done by GraphPad Prism 7(*p<0.05, **p<0.01).

### The expression of human Bax and Bcl-xL is associated with SAM complex

As another series of outer membrane translocases, SAM complex is also worth to study for its necessary role of assisting the functioning of TOM complex. For exploring the relationship between human Bax-induced growth defect and SAM complex. We detect the expression of Sam50, Sam35 and Sam37 with switching on the expression of human Bax and Bcl-xL. The results indicated that the expression of Sam50 and Sam37 was significantly increased by the expression of human Bax individually. When the expression of Bcl-xL was switched on, the expression level of Sam50 even higher. However, the expression of Sam35 did not alter by individual expression of Bax, while it was increased by the expression of both Bax and Bcl-xL (Figure 7).

**Figure 7.**
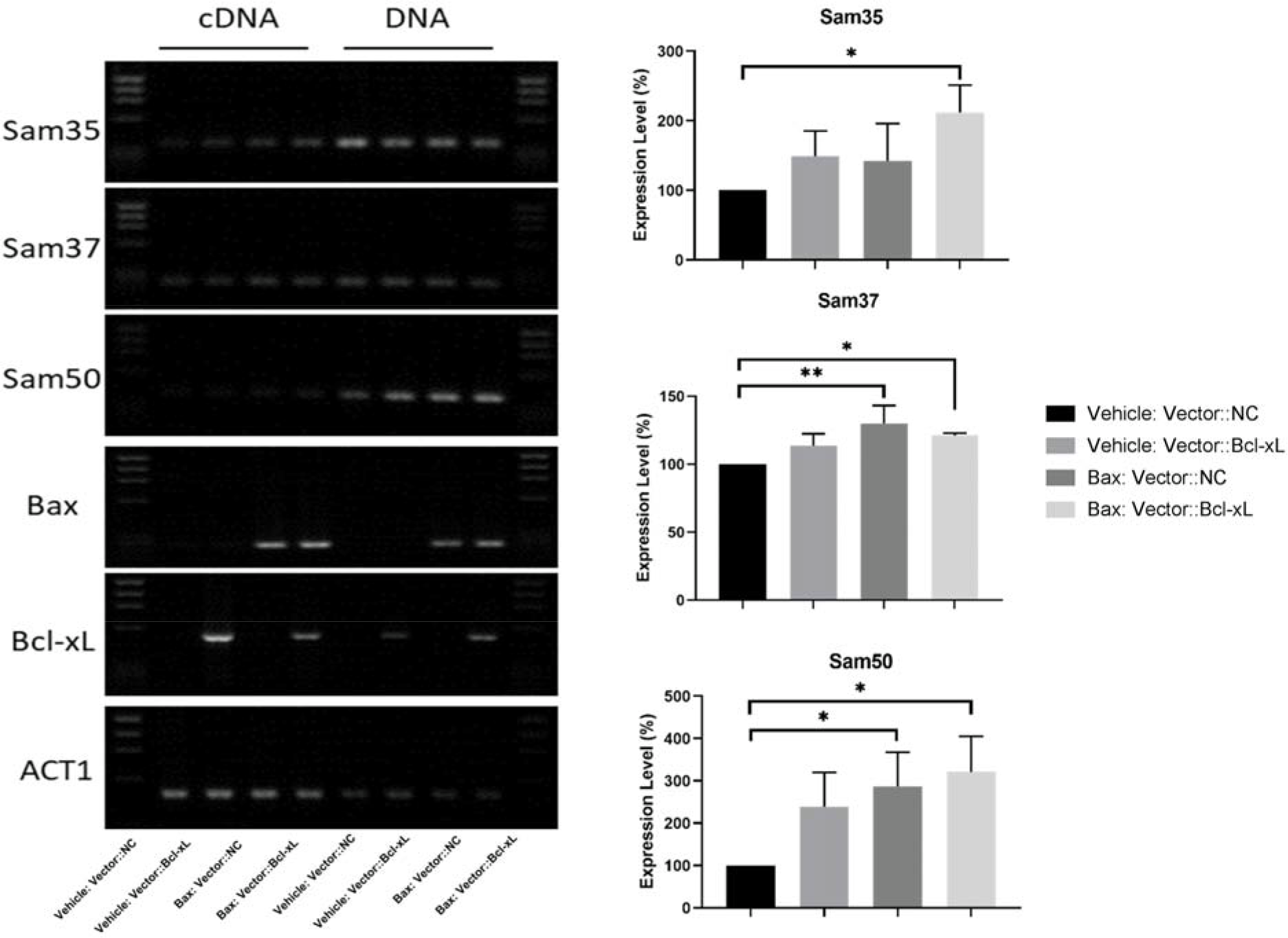
The expression level of the main subunits of SAM complex under the heterologous expression of Bax/Bcl-xL. The images showed that the expression of human Bax and Bcl-xL highly contributes to the increasing expression of Sam37 and Sam50. The expression of Sam35 increased only when switching on the expression of both Bax and Bcl-xL. The PCR products were isolated by 1.5% agarose gel. Gene expression level was measured by the ratios of the grey value of the bands of detected genes and reference gene. The experiment was repeated for three times and the data was presented as mean±SD. Grey value was measured by Image J and One-way ANOVA analysis was done by GraphPad Prism 7(*p<0.05, **p<0.01).

### The expression of human Bax and Bcl-xL have no obvious connection with TIM complex

As the most important translocases of inner membrane, the operation of TIM complex is highly connected with TOM complex during the protein import from outer membrane through the matrix. Although we did not find significant connection between the expression of human Bax/Bcl-xL and TOM complex, it is still necessary to detect the expression level of TIM complex with our Bax/Bcl-xL-integrated yeast system. However, the result revealed that only the expression of Tim50 was decreased slightly by the expression of human Bax. However, this change has no significance. The expression level of Tim23, Tim17 and Tim21 did not alter significantly under the expression of Bax and Bcl-xL (Figure 8).

**Figure 8.**
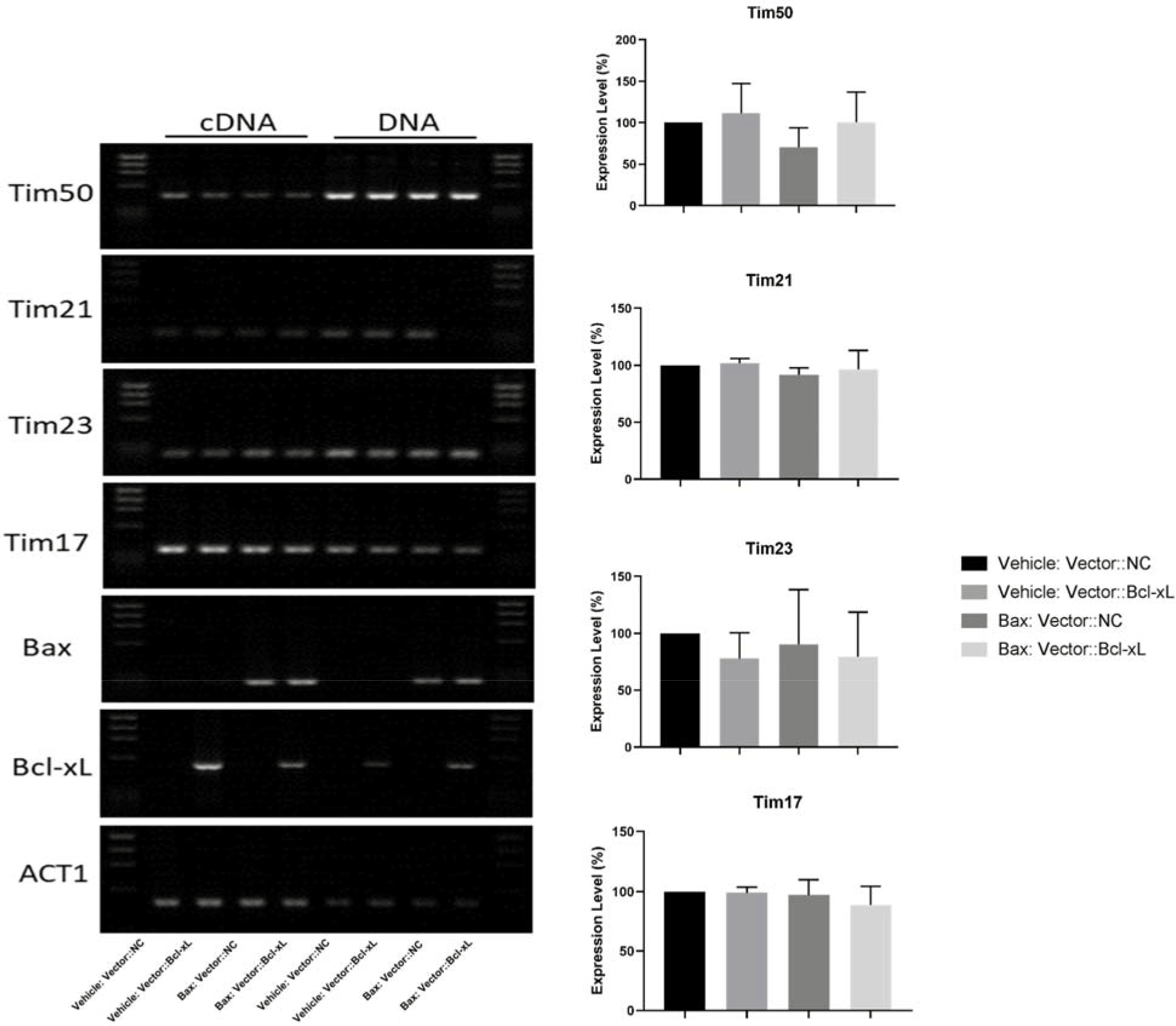
The expression level of four main subunits of TIM complex under the heterologous expression of Bax/Bcl-xL. The images showed that the expression of human Bax is able to inhibit the expression of Tim50. The expression level of another three genes did not change under the expression of Bax/Bcl-xL. The PCR products were isolated by 1.5% agarose gel. Gene expression level was measured by the ratios of the grey value of the bands of detected genes and reference gene. The experiment was repeated for three times and the data was presented as mean±SD. Grey value was measured by Image J and One-way ANOVA analysis was done by GraphPad Prism 7(*p<0.05, **p<0.01).

Based on these findings, it is indicated thatthe human Bax might be able to cause a stress to inhibit the expression of YCA1 and NUC1. Besides, human Bax might cause the growth defect through interacting with SAM complex. Since we have discussed human ECE-1 in introduction, which play an important role in cardiovascular development, cardiac formation and cardiac defect (Hofstra *et al*., 1999); (Takebayashi-Suzuki *et al*, 2000) (Funke-Kaiser *et al*., 2003). For decades, most of the functional studies of the association between ECE-1 and CHD were conducted by using a mammalian cell line or embryos (Funke-Kaiser *et al*., 2003); (Yanagisawa *et al*., 1998). Besides, epidemiological investigation based on ECE-1 polymorphism analysis with the clinical samples of CHD patients was also conducted.

However, it is difficult to study individual isoforms of ECE-1 by using these methods. Furthermore, the molecular basis of CHD remains unclear until now. As we have demonstrated that the pathogenesis of CHD is highly related to apoptosis. Therefore, we focused on two aspects to continue the study: First, to identify the phenotypes of ECE-1 isoforms under Bax-induced growth defect; Second, whether ECE-1 isoforms have ability to affect the expression of candidate genes of mitochondrial outer membrane complex and inner membrane complex.

### The heterologous expression of both Bax and ECE-1 isoforms induces stronger growth defect in yeast, which is inhibited by the expression of Bcl-xL

Four ECE-1 isoforms were transformed into Bax-integrated strains, human Bcl-xL was introduced to the system by doing yeast mating. W303a: Bax/ECE-1 was mated with W303α: pRS404/Gal1p-Bcl-xL-MS, then the three genes were able to express in one diploid cell. The media with galactose is able to switch Gal1 10 promoter on for expressing Bax, Bcl-xL and ECE-1 isoforms, while the cells on glucose media did not trigger the expression. The results indicate that Human ECE-1 did not induce growth defect alone, interestingly, when expressing with Bax, these isoforms seem to act as an enhancer of Bax-induced growth defect. Particularly, ECE-1c has the strongest effect. Interestingly, the expression of Bcl-xL was able to protect cells from Bax-induced cell death and the effect enhanced by ECE-1 isoforms compared with the controls. However, the strain that express both Bax and ECE-1b was not sensitive to the expression of Bcl-xL (Figure 9).

**Figure 9.**
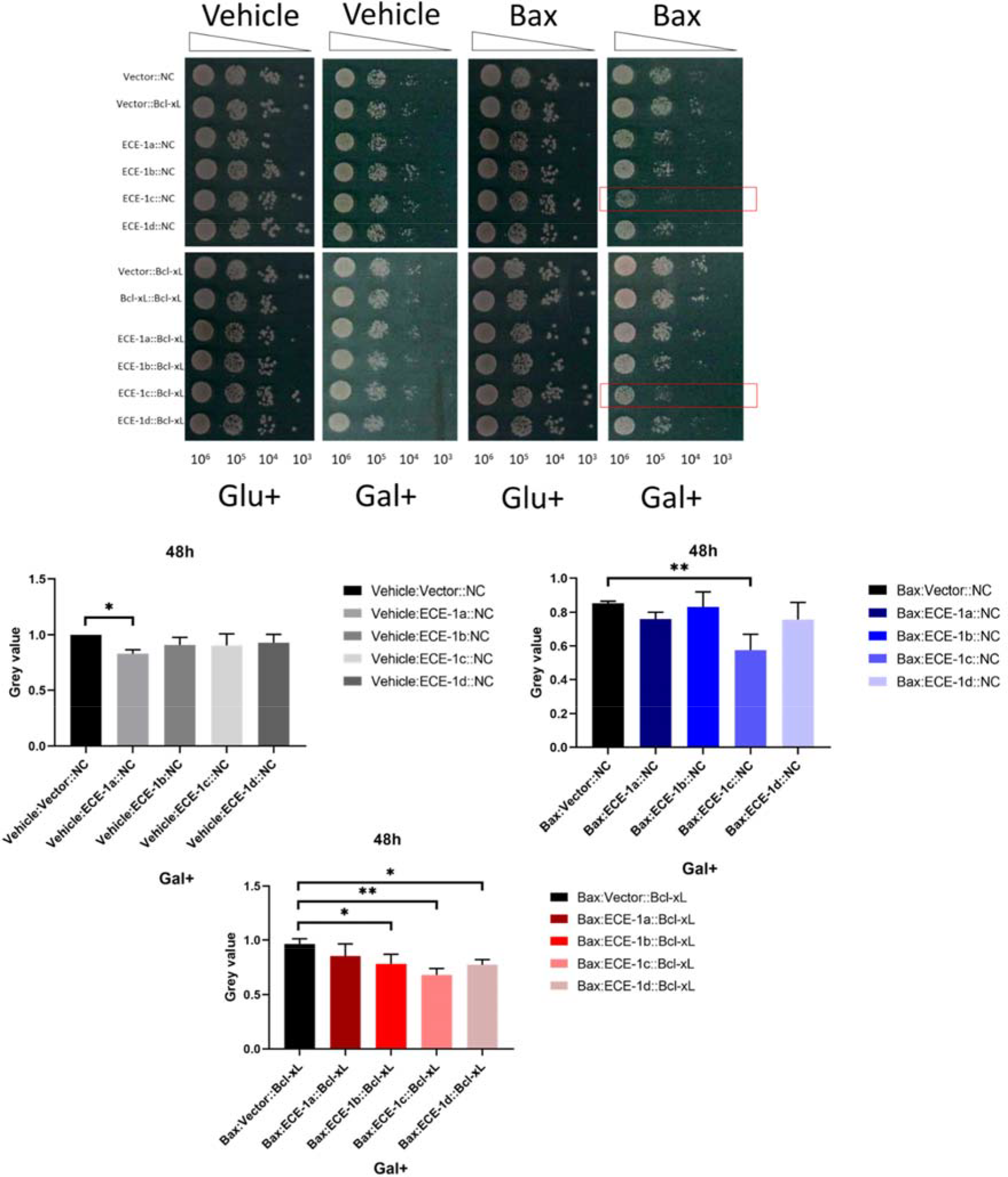
The representative image of spot assay for the yeast diploids expressing Bax and ECE-1 isoforms as well as pRS404/Bcl-xL on SD/SG-LEU-URA-TRP plates. The plates were incubated in in 30 °C for 48h. The harvested cells were processed with 10-fold dilution, which ranging from 10^6^ cells to 10^3^ cells. The experiments were repeated for 3 times and spots of undiluted group were chosen and analysed with Image J for measuring grey values. The mean of grey values was calculated as well as the SDs compared with control group (the first bar of each graph). Ordinary one-way ANOVA was conducted by Graphpad Prism 7 (*p<0.05, **p<0.01).

### The growth rate of yeast cells was inhibited under the heterologous expression of Bax, ECE-1 isoforms and Bcl-xL

Following the results from the spot assay, the growth curves of the diploids containing pRS404/Gal1p-Bcl-xL-MS were recorded. These strains were cultivated automatically on a SG-URA-LEU-TRP drop-out agar plate, the size of which is suitable for the ROTOR HDA system. The incubation was conducted in microplate reader at 30°C for 65h. The growth curves were recorded by microplate reader and export from the PHENOS software (Barton et al, 2018). The curves exactly matched with the previous results in spot test: The growth rate of yeast was inhibited with the expression of both Bax and ECE-1 isoforms and Bcl-xL was able to protect the cells slightly. Four ECE-1 isoforms have no effect on inducing cell death individually. However, they enhance the effect of Bax and inhibit the growth of cells in different levels. When Bcl-xL was introduced into the strains, the cells begin to survive from this growth defect. Interestingly, the Bax-induced cell death, which enhanced by ECE-1b seem to be not sensitive to the expression of Bcl-xL compared with others as shown in spot assay (Figure 10).

**Figure 10.**
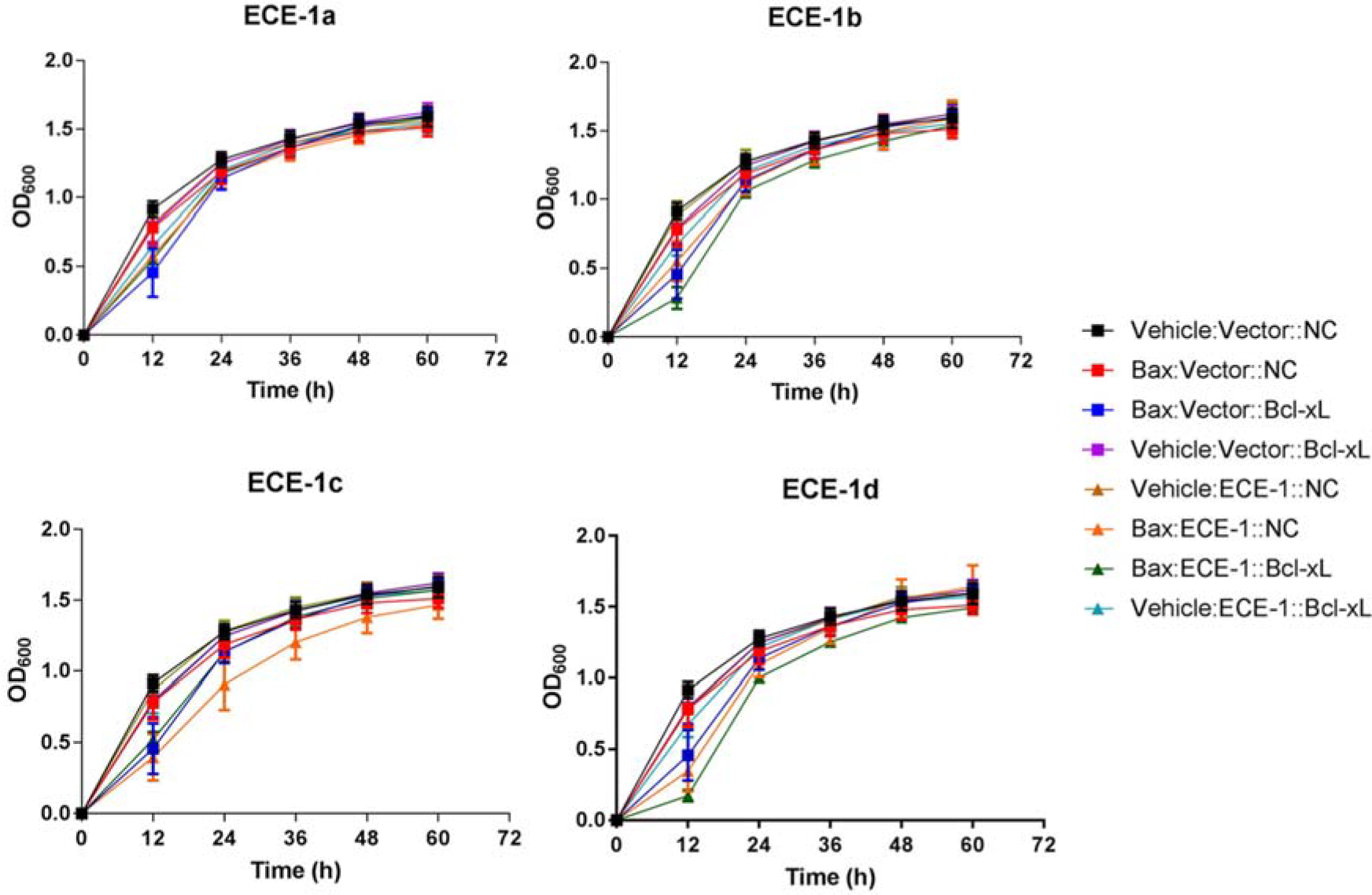
The growth curves of the diploid yeast cells with the expression of Bax, ECE-1 isoforms and Bcl-xL. The strains were spotted on SG-LEU-URA-TRP drop out agar plates by Singer Rotor, then incubated and analysed by microplate reader at 30 °C. The OD_600_ was measured constantly during 65h. The data was exported by using PHENOS and we choose five OD_600_ values of each strain for every 12h, the curves were made by Graphpad Prism 7.

### ECE-1c enhanced Bax-induced growth defect inhibited the expression level of NUC1 but has no influence on the expression of YCA1 and BI-1

It is obvious that individual ECE-1 isoforms have no influence on causing yeast growth defect. However, all the isoforms are able to induce growth defect under the expression of Bax, particularly, ECE-1c is proved to be the most sensitive one. As a most well-studied isoforms of ECE-1, ECE-1c has been confirmed to be the most widely located and expressed in almost all tissues (Lindenau *et al*, 2006). Besides, ECE-1c is the only one located mainly on plasma membrane among all the ECE-1 isoforms (Schweizer *et al*., 1997). Therefore, we suppose that ECE-1c is the most possible one to have an interaction with the mitochondria. Therefore, ECE- 1c will be involved in the further study for investigating the mRNA level of mitochondrial genes that have been assessed in Bax/Bcl-xL yeast system.

In order to investigate the role of ECE-1c in Bax-induced growth defect, the expression level of YCA1, NUC1 and BI-1 was assessed with adding the heterologous expression of ECE-1c based on the results in chapter 4. The results showed that the individual ECE-1c did not altered the expression level of any of the three genes, however, the expression level of NUC1 decreased sharply under the expression of both Bax and ECE-1c, while the expression of Bcl-xL made it recovered to the normal level (Figure 11).

**Figure 11.**
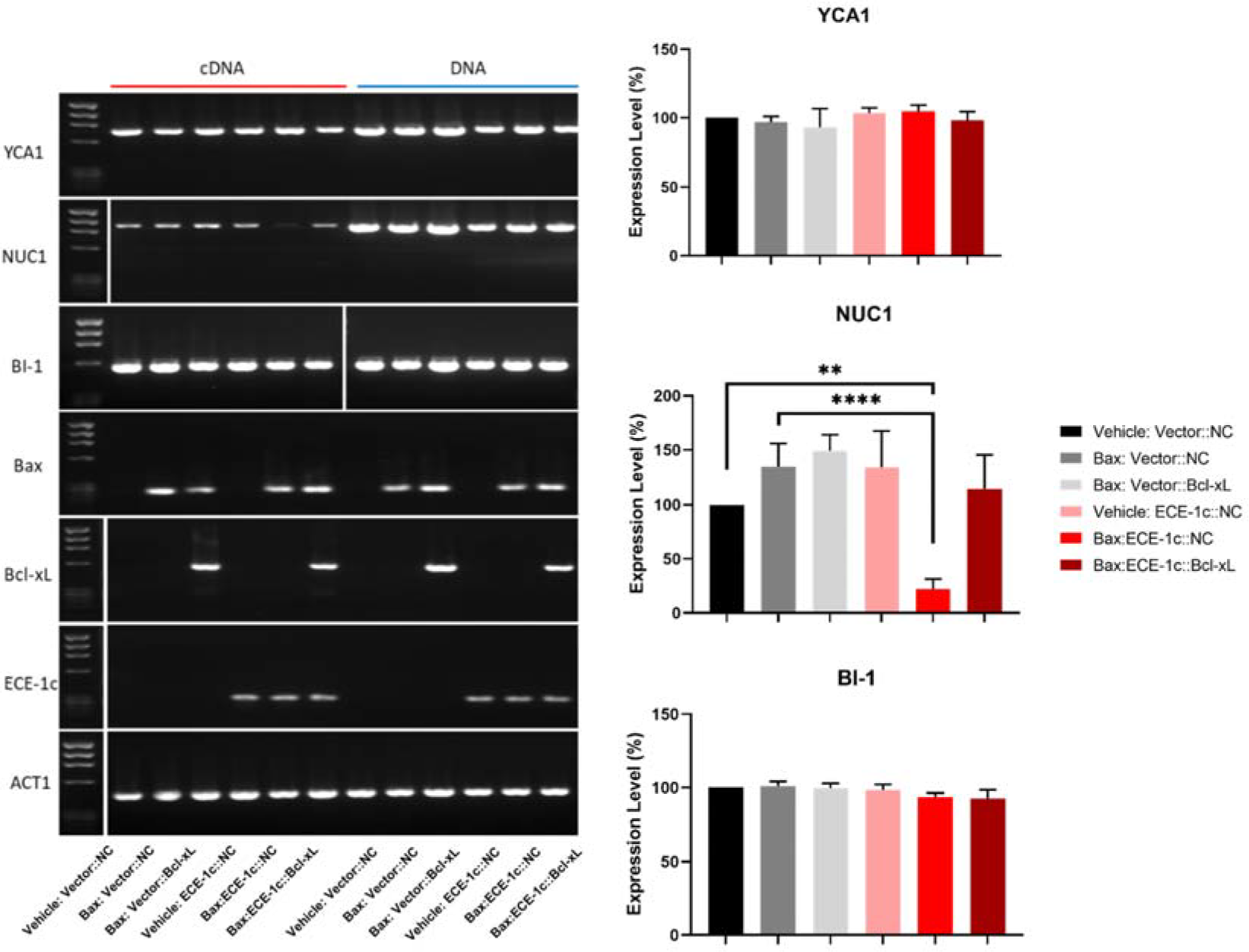
The expression level of three representative yeast apoptotic molecules under the ECE- 1c enhanced Bax-induced growth defect. The image indicate that Bax-induced growth defect enhanced by ECE-1c inhibit the expression level of NUC1 but have no effect on YCA1 and BI-1. The PCR products were isolated by 1.5% agarose gel. Gene expression level was measured by the ratios of the grey value of the bands of detected genes and reference gene. The experiment was repeated for three times and the data was presented as mean±SD. Grey value was measured by Image J and One-way ANOVA analysis was done by GraphPad Prism 7 (**p<0.01, ***p<0.001).

### The expression of receptors of TOM complex is decreased by ECE-1c enhanced Bax-induced growth defect

It has been investigated that heterologous expression of Bax/Bcl-xL did not affect the mRNA level of the six subunits of TOM complex but increased the expression of Sam37 and Sam50. Interestingly, a previous study also measured the expression of several subunits of TOM complex and SAM complex by western-blotting analysis under the heterologous expression of human Bax, it has reported that both TOM and SAM complex are not necessary for the interaction between Bax and mitochondria (Sanjuán Szklarz *et al*, 2007). In addition, it is convinced that ECE-1c have a powerful effect on enhancing Bax-induced growth defect according to the results of spot assay and cell viability assay in this chapter. Therefore, we decided to assess whether TOM complex will be affected under the ECE-1c enhanced Bax-induced growth defect. According to the results, the individual ECE-1c did not alter the expression level of any of the Tom subunits, however, the expression level of two receptors Tom70 and Tom22 was significantly decreased under Bax-induced growth defect enhanced by ECE-1c. The expression level of channel-forming unit Tom40 was also inhibited by ECE-1c enhanced Bax-induced growth defect, while another three subunits Tom20, Tom5 and Tom6 were not influenced (Figure 12) in a same condition.

**Figure 12.**
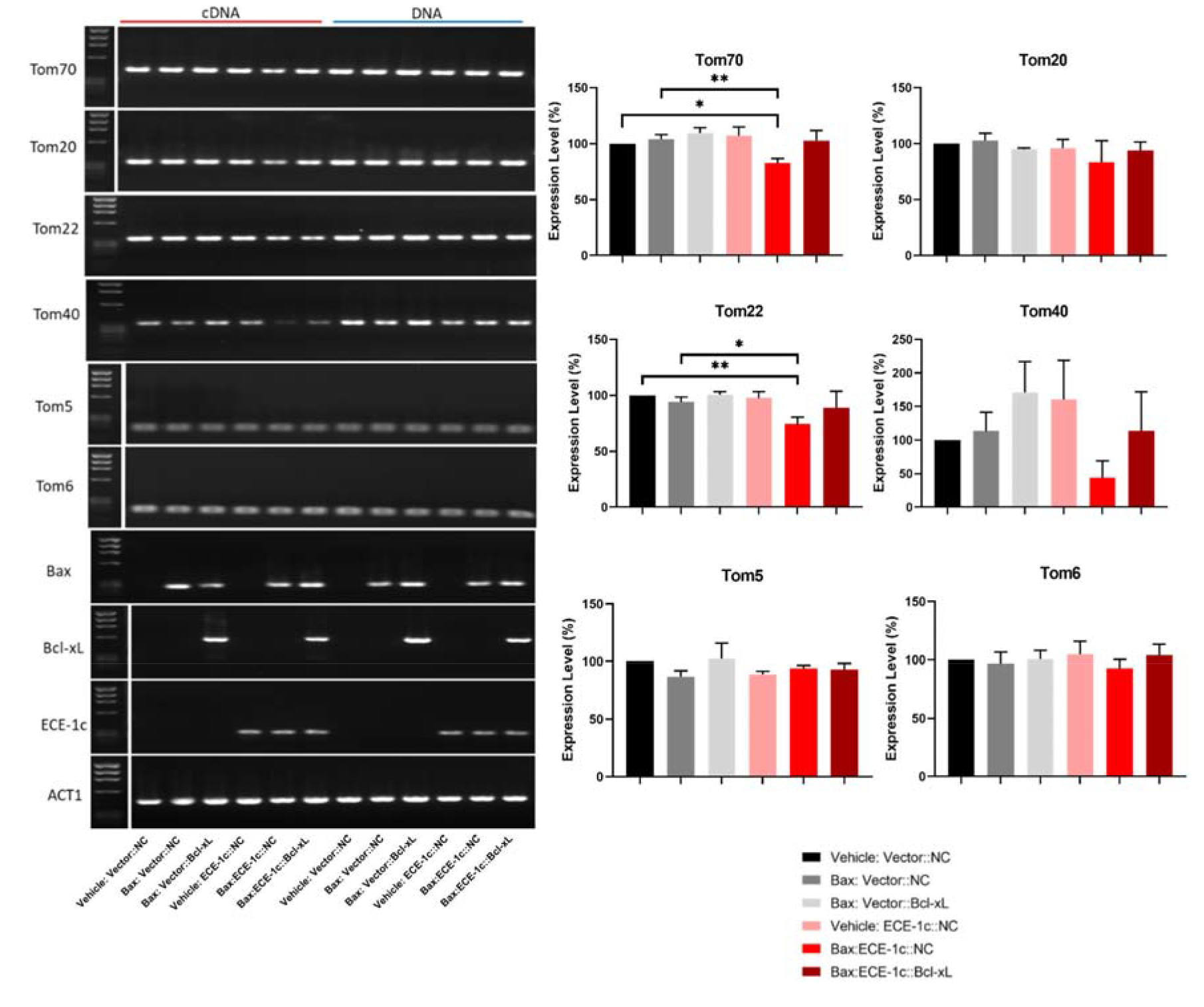
The expression level of representative subunits of TOM complex under the ECE-1c enhanced Bax-induced growth defect. The image showed the individual ECE-1c have no effect on TOM complex. A decreasing expression level of Tom70 and Tom22 caused by ECE-1c enhanced Bax-induced growth defect was observed, which is inhibited by Bcl-xL and return to an original level. The PCR products were isolated by 1.5% agarose gel. Gene expression level was measured by the ratios of the grey value of the bands of detected genes and reference gene. The experiment was repeated for three times and the data was presented as mean±SD. Grey value was measured by Image J and One-way ANOVA analysis was done by GraphPad Prism 7 (*p<0.05, **p<0.01).

### ECE-1c enhanced Bax-induced growth defect has an effect on SAM complex in contrast to individual Bax-induced growth defect

It has been demonstrated that individual Bax-induced growth defect might be associated with SAM complex by promoting the expression level of three subunits (Sam35, Sam37 and Sam50). However, it is found that an enhanced Bax-induced growth defect by ECE-1c inhibit the expression level of Tom receptors strongly. Therefore, as SAM complex is another outer membrane machinery that correlated with TOM complex closely, it is significant to investigate the expression level of the three subunits of SAM complex under the ECE-1c enhanced Bax-induced growth defect. The results indicated that individual ECE-1c have no effect on SAM complex. Interestingly, the expression level of Sam35 and Sam50 decreased dramatically under ECE-1c enhanced Bax-induced growth defect, while the expression of Bcl-xL is able to inhibited this synergistic effect. The expression level of Sam37 remained as usual (Figure 13).

**Figure 13.**
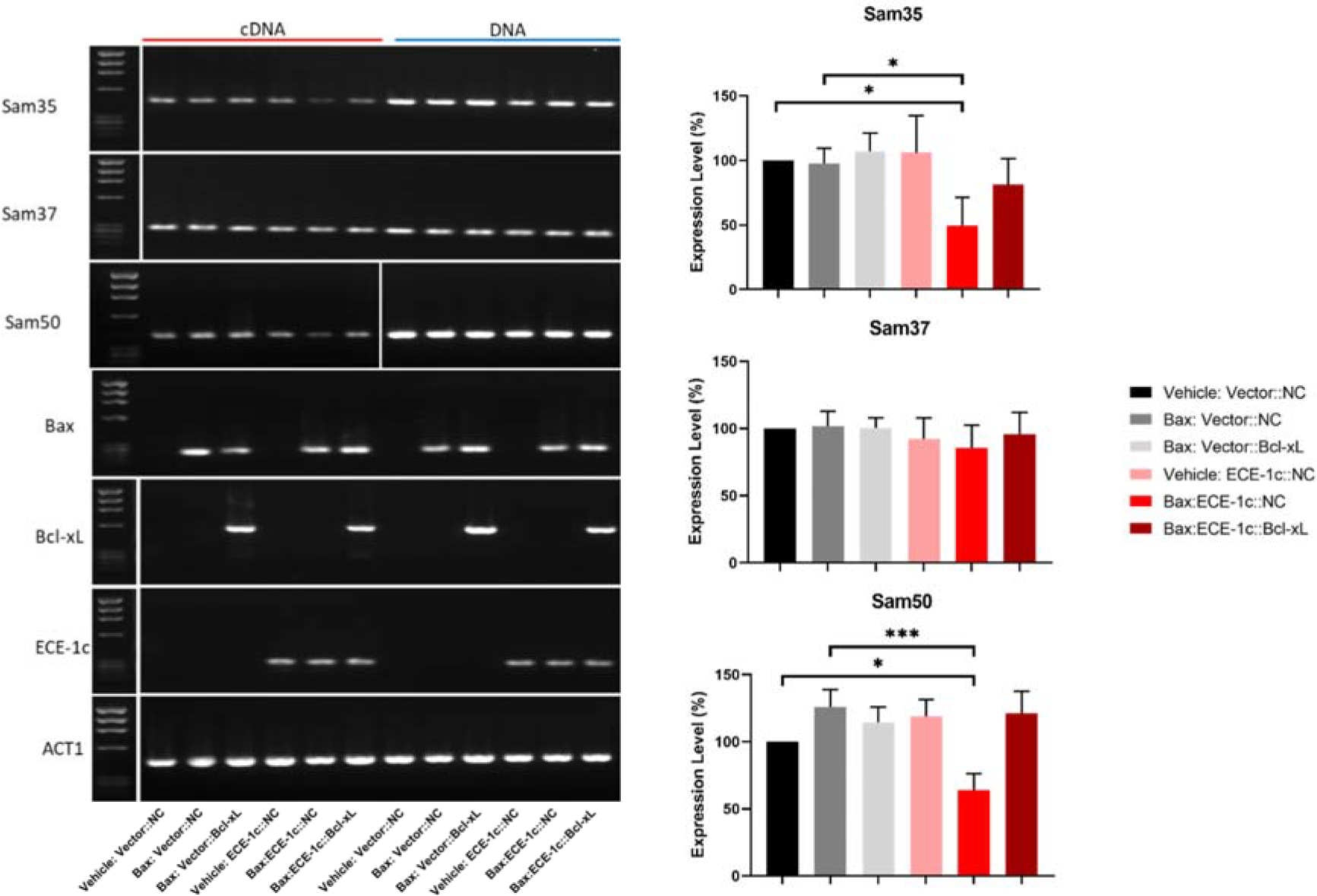
The expression level of the main subunits of SAM complex under the ECE-1c enhanced Bax-induced growth defect. The image indicated the expression level of Sam35 and Sam50 was decreased by ECE-1 enhanced Bax-induced growth defect. Individual ECE-1c showed no influence on SAM complex. The expression of Sam37 was not altered. The PCR products were isolated by 1.5% agarose gel. Gene expression level was measured by the ratios of the grey value of the bands of detected genes and reference gene. The experiment was repeated for three times and the data was presented as mean±SD. Grey value was measured by Image J and One-way ANOVA analysis was done by GraphPad Prism 7 (*p<0.05, ***p<0.001).

### The expression of Tim complex is not altered by both individual ECE-1c and ECE-1c enhanced Bax-induced growth defect

According to the results from (Figure 12) and (Figure 13), it is concluded that the enhancement of ECE-1c in Bax-induced growth defect contribute a significant inhibition to the expression of outer membrane translocases, which is totally different from the situation under individual Bax-induced growth defect. Hence, it is interesting to explore whether this synergistic effect of Bax and ECE-1c is able to affect the stability of inner membrane through TIM complex. Although the results in (Figure 8) illustrated that TIM complex of inner membrane was not affected by Bax-induced growth defect, we suppose the ECE-1c enhanced Bax-induced growth defect would be different. Therefore, the expression level of receptors Tim50, Tim21, the channel-forming units Tim23 and Tim17 were assessed with involving ECE-1c. However, the result was exactly same as it appeared in (Figure 8): The expression level of all four Tim subunits did not change (Figure 14).

**Figure 14.**
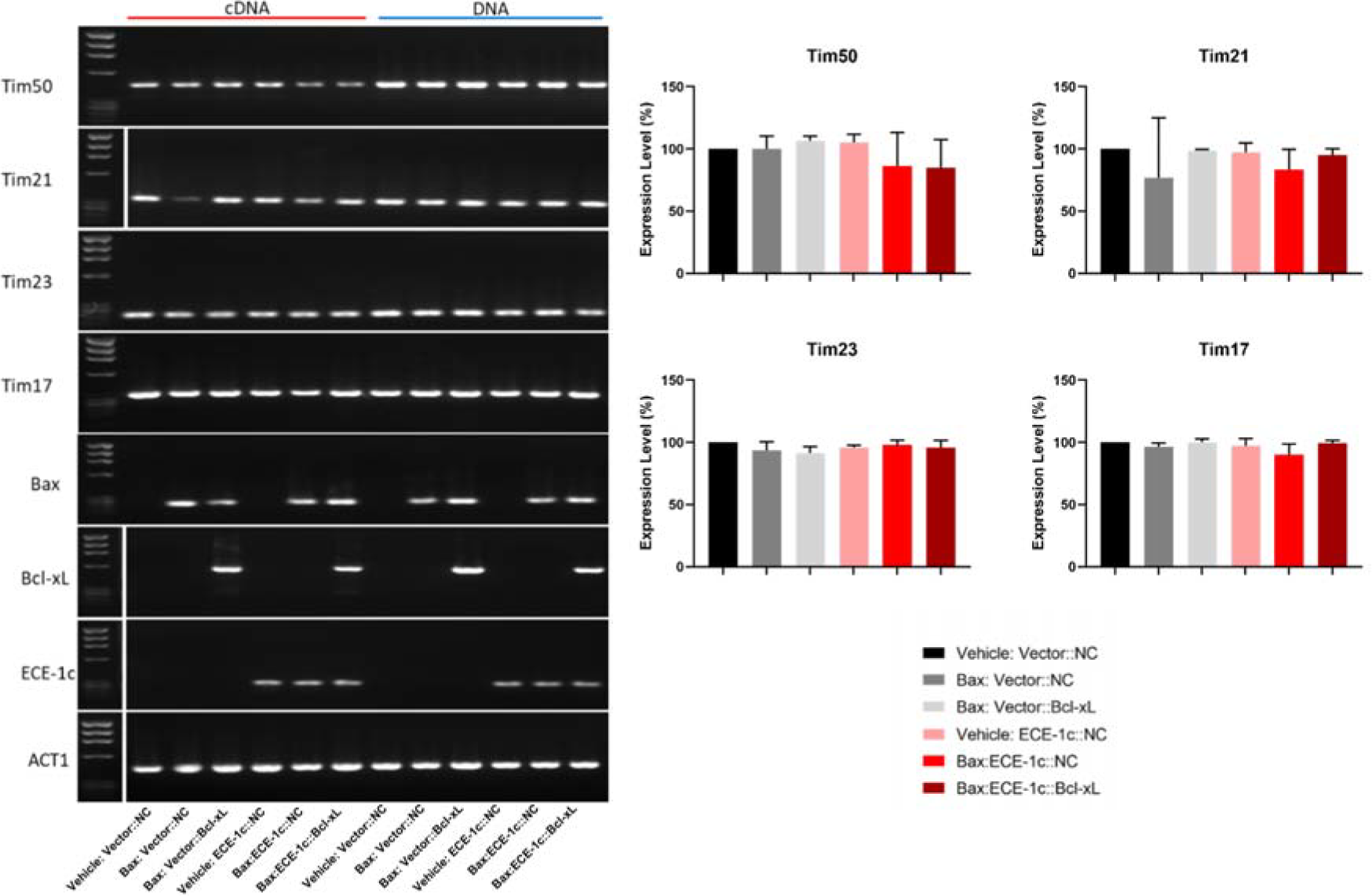
The expression level of four main subunits of TIM complex under the ECE-1c enhanced Bax-induced growth defect. The image indicated that both ECE-1c and ECE-1c enhanced Bax-induced growth defect have no influence on TIM complex. The PCR products were isolated by 1.5% agarose gel. Gene expression level was measured by the ratios of the grey value of the bands of detected genes and reference gene. The experiment was repeated for three times and the data was presented as mean±SD. Grey value was measured by Image J and One-way ANOVA analysis was done by GraphPad Prism 7.

## Discussion

In fact, to mimic human Bax-induced apoptosis in yeast is not a new technique. There is one group have conducted the humanization of mammalian Bax and Bcl-xL in a *S. cerevisiae* strain named WCG4 (Ligr *et al*., 1998), Bax and Bcl-xL was inserted into plasmid pSD10.a respectively for transformation, another two plasmids named pRS315 and pRS316 were also transformed into WCG4 as control vectors. Therefore, it is convinced that Bax/Bcl-xL expression system of yeast is available for studying the function of human genes that correlated with mitochondrial pathway. In our study, a new yeast system for the mimicking of Bax-induced apoptosis was generated by involved in human Bcl-xL with two patterns: The original insert of Bcl-xL was digested from a 2-micron plasmid pRS426, which was ligated to another 2-micron plamid of pRS424 and an integrative plasmid of pRS404. Why two kinds of vectors were chosen in this step? The pRS424 is a high copy number plasmid, which is able to replicate independently from yeast genome due to its 2µ circle replication region. This characteristic might also increase the copy number of Bcl-xL and highlights its anti-apoptotic effect during the apoptosis mimicking in yeast. While the pRS404, which is a single copy plasmid for its lack of a replication origin. Therefore, the replication of pRS404 have to depend on its integration into yeast genome. This might not be an advantage for observing the phenotype of Bcl-xL in the yeast system due to the low copy number of pRS404, however, this characteristic brings the stability for the expression of Bcl-xL, because genome DNA is much more stable and has less chance to lose from the yeast cells. However, there were only the strains carrying pRS404 presented changes of phenotypes, the strains carrying pRS424 have no differences in phenotypes. This might indicate that multi-copied Bax/Bcl-xL results are different from single-copied Bax/Bcl-xL results in mRNA or protein level. Therefore, it is necessary to think about detecting the expression in mRNA or protein level in further study by qPCR and Western-blotting. Fortunately, the new humanized yeast system has been established successfully and ready for using in further gene function analysis.

Based on the phenotypes of human Bax/Bcl-xL in yeast through the spot assay, we have investigated the correlation between human Bax-induced growth defect and the mRNA levels of several genes of yeast mitochondria with the strains containing pRS404. YCA1, NUC1 and BI-1 were tested as the first experiment. YCA1 and NUC1 are highly associated with mitochondrial pathway in yeast for inducing apoptosis (Gowsalya *et al*, 2019). The BI-1 is a important regulator for preventing the cell death induced by pro-apoptotic protein of Bcl-2 family (Kawai *et al*, 1999). We supposed that the expression level of these three genes would increase, however, the results were completely different from our expectation: The expression level of YCA1 and NUC1 decreased, while BI-1 has no significant change under Bax-induced growth defect. The possible reason is that heterologous expression of human Bax might not induce apoptosis depend on mitochondrial pathway. A previous study has measured the correlation between the expression of YCA1 and human Bax using immunoblot analysis, it has been reported that the activation of YCA1 is not required for Bax toxicity (Guscetti *et al*, 2005). Therefore, it is resonable to justify that human Bax-induced growth defect might not able to cause apoptosis but only a kind of stress, which is independent from the activation of YCA1, based on this, it is understandable that the expression of BI-1 is not altered by Bax. However, we are not able to clarify whether apoptosis contribute to the Bax-induced growth defect by gene expression assay alone. The flow cytometry after staining the yeast cells by Annexin V and PI will be a direct method for recognizing apoptotic cells, the morphological changes of which are able to be analysed under fluorescence microscope. For these assays for measuring apoptosis directly, pRS404 strains would be the better choice than pRS424 strains because of the clearer phenotypes have been identified in spot assay, it is not sure whether the growth of pRS424 strains were inhibited according to the phenotypes shown in spot assay, hence there might be difficult to find apoptotic cells or other morphological changes.

The mitochondria protein import translocases play the critical role in recognizing and translocating the precursor proteins or targeting signals (Wiedemann *et al*., 2004). Our study tested several subunits of TOM complex, SAM complex and TIM complex. It has been shown that only the expression of Sam35, Sam37 and Sam50 increased under Bax-induced growth defect while no significant change is found in other genes of TOM and TIM complex. Interestingly, the co-expression of Bax and Bcl-xL increased the mRNA level of three subunits of SAM complex stronger than individual Bax. Therefore, we speculate that human Bax cause the growth defect through the interaction with SAM complex of mitochondrial outer membrane, since SAM complex is responsible for the process of β-barrel proteins, the expression of Bax and Bcl-xL might have a synergistic effect on promoting the formation and insertion of β- barrels. However, the results of mRNA levels were not able to clarify the relationship between heterologous expression of Bax and these mitochondrial genes thoroughly. There are two aspects should be included in the improvement of gene expression assay: First, to measure the protein levels of these mitochondrial genes using western-blotting analysis, which have shown the positive results in mRNA levels; Second, to knockout the mitochondrial genes respectively that have shown the positive results in mRNA levels, then to confirm the protein levels of Bax and Bcl-xL in both knockout strains and non-knockout strains, the results of which would be able to explain whether these mitochondrial genes are required for Bax-induced growth defect in yeast. In addition, it would be meaningful to carry out the improved gene expression assay using both pRS424 and pRS404 strains, because we are not able to know the protein levels according to the phenotypes of spot assay.

The function of human ECE-1 isoforms have been identified preliminaryly by evaluating the yeast growth defect through spot assay and growth curves recording. It is confirmed that individual ECE-1 isoforms cause no growth defect in yeast, however, all of these isoforms are able to enhance the growth defect caused by Bax. The 65h growth curves under the heterologous expression indicated that yeast cells increased rapidly during first 12h and stay in exponential phase until 24h, the growth rate of yeast cells is dramatically inhibited under the co-expression of Bax and ECE-1 isoforms during exponential phase, ECE-1c caused the strongest inhibitiaon. The growth curves started to turn into the end of exponential phase after 24h and woud go into the stationary phase soon. This means the numbers of yeast cells were approximately close to the maximum that could be harvested at this time. Therefore, we chose 24h as the time for extracting DNA and RNA and carried out the heterologous expression in the further investigation of the role of ECE-1c in affecting the expression of candidate genes of mitochondrial translocases.

The expression of individual ECE-1c is unable to affect the expression of these mitochondrial genes. After the expression level of these candidate genes was measured under ECE-1c enhanced Bax-induced growth defect, only the expression level of NUC1, two receptors of TOM complex (Tom70, Tom22) and Sam35, Sam50 decreased significantly. In fact, three different views about the interaction between Bax and mitochondria have been reported: 1) Bax-induced apoptosis is actually a toxicity to mitochondria, which results in a damage on outer membrane (Priault *et al*, 1999); 2) Bax has an ability of channel-forming on outer membrane, the interaction between Bax and mitochondria is independent from other endogenous proteins (Kuwana *et al*, 2002); 3) Other outer membrane translocases are essential to the channel formation of Bax (Pavlov *et al*., 2001). Hence, It is convinced that the growth defect caused by co-expression of Bax and ECE-1c is highly correlated with a series of mitochondrial genes. It is known that NUC1 is released from mitochondria as an endogenous pro-apoptotic molecule, therefore, the inhibition caused by Bax and ECE-1c means the release of NUC1 might be blocked due to the permeabilization has been changed by a disruption on outer membrane. For the decreasing expression level of the Tom70, Tom22 under the co-expression of Bax and ECE-1c, we speculte that Bax is able to form a mitochondrial apoptosis-induced channel (the MAC that have been discussed in the introduction) across the outer membrane with the assisstant of ECE-1c, the permeabilization of mitochondria might be altered. Therefore, the function of main receptor and secondary receptor of outer membrane, which are Tom70 and Tom22 were likely to be disturbed or suppressed when the Bax interacts with mitochondrial outer membrane. The decreasing of the expression of Sam35 and Sam50 means ECE-1c enhanced Bax-induced growth defect may affect the formation or function of β-barrel protein due to the folding of β-barrel accomplished by Sam50-Sam35 complex. However, the results of TIM complex indicated that ECE-1c enhanced Bax-induced growth defect did not affect the inner membrane. Interestingly, all the positive results involving the expression of ECE-1c were completely different from the results of the expression of individual Bax. Therefore, it is possible that Bax-induced growth defect is able to alter the permeabilization of mitochondrial outer membrane through interacting with the receptors of TOM and SAM complex with the assisstant of ECE-1c. However, these results were insufficient to explain whether the synergistic effect of Bax and ECE-1c is apoptosis, necrosis or a kind of stress. As the gene expression assay is an indirect method to study apoptosis, it is necessary to follow the study by measuring the apoptosis directly. Moreover, the gene expression assay of mRNA levels in our study is also inadequate to clarify whether the mitochondrial genes are required during the ECE-1c enhanced Bax-induced growth defect. For following this question, it is critical to measure the protein expression levels of ECE-1c, Bax and Bcl-xL after knocking out these mitochondrial genes respectively.

For decades, most of the studies of CHD related to ECE-1 remain in the level of analysing genotypes among the clinical samples of CHD (Wang *et al*., 2012b), or the immunoblotting analysis based on a mammlian model (Yanagisawa *et al*., 1998). However, it is impossible to study individual isoforms of ECE-1 with a mammlian system, the advantage of a humanized yeast system is to simplify this problem. In the future, this humanized yeast system might be used to test other interesting genotypes (e.g. mutations of *ACTC1, GATA4, GATA6*) (Augiere *et al*., 2015) (Wang *et al*, 2013) (Wang *et al*, 2012a) related to CHD directly in the for the potential function of regulating apoptosis or affecting mitochondria. Moreover, the humanized yeast system hope to be used for screening cDNA libraries of human heart tissues with CHD and without CHD for potential anti-apoptotic regulators: First, to express two cDNA libraries individually in Bax-integrated strains; Second, to screen for yeast colonies, which potentially contain an anti-apoptotic gene and compare the differences in the expression of their anti-apoptotic proteins between two cDNA libraries have on the screening; Finally, to identify these phenotypes related to CHD from yeast to mammalian cells.

Although our findings in yeast did not reveal the possible function of ECE-1 in CHD directly, it provides two useful evidence: 1) ECE-1 isoforms have an ability to contribute to the apoptosis caused by Bax; 2) ECE-1c is highly correlated with the stability of mitochondrial outer membrane during Bax-induced apoptosis. These might be new perspectives for the treatment of CHD and perhaps ECE-1 is able to act as a biomarker of the prognosis of CHD.

## Materials and Methods

### Strains, media and vector plasmids used

DH5α (F– Φ80lacZΔM15 Δ (lacZYA-argF) U169 recA1 endA1 hsdR17 (rK–, mK+) phoA supE44 λ– thi-1 gyrA96 relA1) (Invitrogen, Paisley, UK) was used for the routine cloning work. W303 MATa (leu2-3,112 trp1-1 can1-100 ura3-1 ade2-1 his3-11, 15) and W303 MATα (leu2-3,112 trp1-1 can1-100 ura3-1 ade2-1 his3-11, 15) were used to express the new constructed plasmids.

Luria-Bertani (LB) broth and LB solid media were used for bacterial culture and DH5α transformation (The LB media for transformation is consisted of 1% w: v Tryptone, 0.5% w: v Yeast Extract, 1% w: v NaCl, 100µg/ml Ampicillin, 1.5% w: v Agar if spread plates). SOC media was used for DH5α transformation (2g Bacto Tryptone, 0.5g Bacto Yeast Extract, 0.2ml 5M NaCl, 0.25ml 1M KCl, 1ml 1M MgCl_2_, 1ml MgSO_4_, 2ml 1M Glucose. Yeast extract-peptone-dextrose (YPD) was used for yeast cell culture (4g Yeast Extract, 8g Bacto Peptone, 8g D-Glucose, 4ml 0.5% Adenine Solution, 8g Agar if spread plates) (Per 400ml). SD-TRP media was used for yeast transformation (2.7g Yeast nitrogen w/o amino acid, 8g D-Glucose, 0.35g TRP drop-out powder, 2.5ml 1% leucine and 1.2ml 1% Lysine solution, pH6-6.5) (Per 400ml). SD/SG-LEU-TRP SD/SG-LEU-URA and SD/SG-LEU-URA-TRP media were used for spot assay (2.7g Yeast nitrogen w/o amino acid, 8g D-Glucose/ or 32ml 25% Galactose solution, 0.35g -LEU-TRP/-LEU-URA/ -LEU-URA-TRP drop-out powder, 1.2ml 1% Lysine solution, 8g Bacto Agar, pH6-6.5) (Per 400ml). The drop-out powder was mentioned above is a mixture of variety of amino acid powder for the growth of normal yeast cells. It includes: 0.8g Arginine, 0.8g Adenine, 4g Aspartic acid, 0.8g Histidine, 0.8g Leucine, 1.2g Lysine, 0.8g Methionine, 2g Phenylalanine, 8g Threonine, 0.8g Tryptophan, 1.2g Tyrosine, 0.8g Uracil. Before making this powder, we need to make sure not add the amino acid that is shown on selectable markers.

pRS426 containing the Gal1 promoter, C-terminal Myc-tag and SUC2 terminator (MS) and the CDS sequence of Bcl-xL is a 2-micron (2µ) plasmid. pRS424 and pRS404 (Chee & Haase, 2012) both contain a selectable marker of TRP1, which were used in the cloning experiments as vectors for new constructions. The former one is a 2-micron plasmid while the latter one is an integrating plasmid (YIp), which needs to be integrated into the host chromosome for expression.

The coding sequence (CDS) of human Bax was integrated into another integrative plasmid pRS405, which was introduced to W303 strain during the previous work of lab. The sequence of four ECE-1 isoforms were cloned from human HEK293 cells by Reverse-transcription PCR and were also inserted into a 2-micron plasmid previously, which is pRS426 (Lu & Willars, 2019). Both the new constructed pRS405 and pRS426 were confirmed by sequencing.

### Primers used and the generation of Bax/Bcl-xL-integrated system

The primers used in this study were designed for the confirmation of full length of 4 new constructed plasmids. M13 primers, specific primers named Gal575 and SUC75 were used to confirm the insert DNA of new constructs (Table 1). The primers designed for detecting the expression level of representative mitochondrial genes were shown as

Table *2*.The establishment of the yeast system begins with pRS426/Gal1p-MS and pRS426/Gal1p-Bcl-xL-MS, which were constructed previously by another colleague in our lab and it worked well in W303 strains. pRS426 is a 2-micron plasmid (high-copy number) with a selectable marker of URA3, two inserts were moved from pRS426 by the digestion with XhoI and SacI then ligated to pRS424 and pRS404 respectively. pRS424 is another 2-micron plasmid while pRS404 is an integrative plasmid, they share the same selectable marker of TRP1. One of the inserts contain Gal1 10 promoter, C-terminal Myc-tag, SUC Terminator and a stop codon after Myc, another is exactly same except a coding DNA sequence (CDS) of human Bcl-xL between Gal1 10 promoter and Myc. After the ligation, four new constructs were generated (Figure 15) (Figure 16).

**Figure 15.**
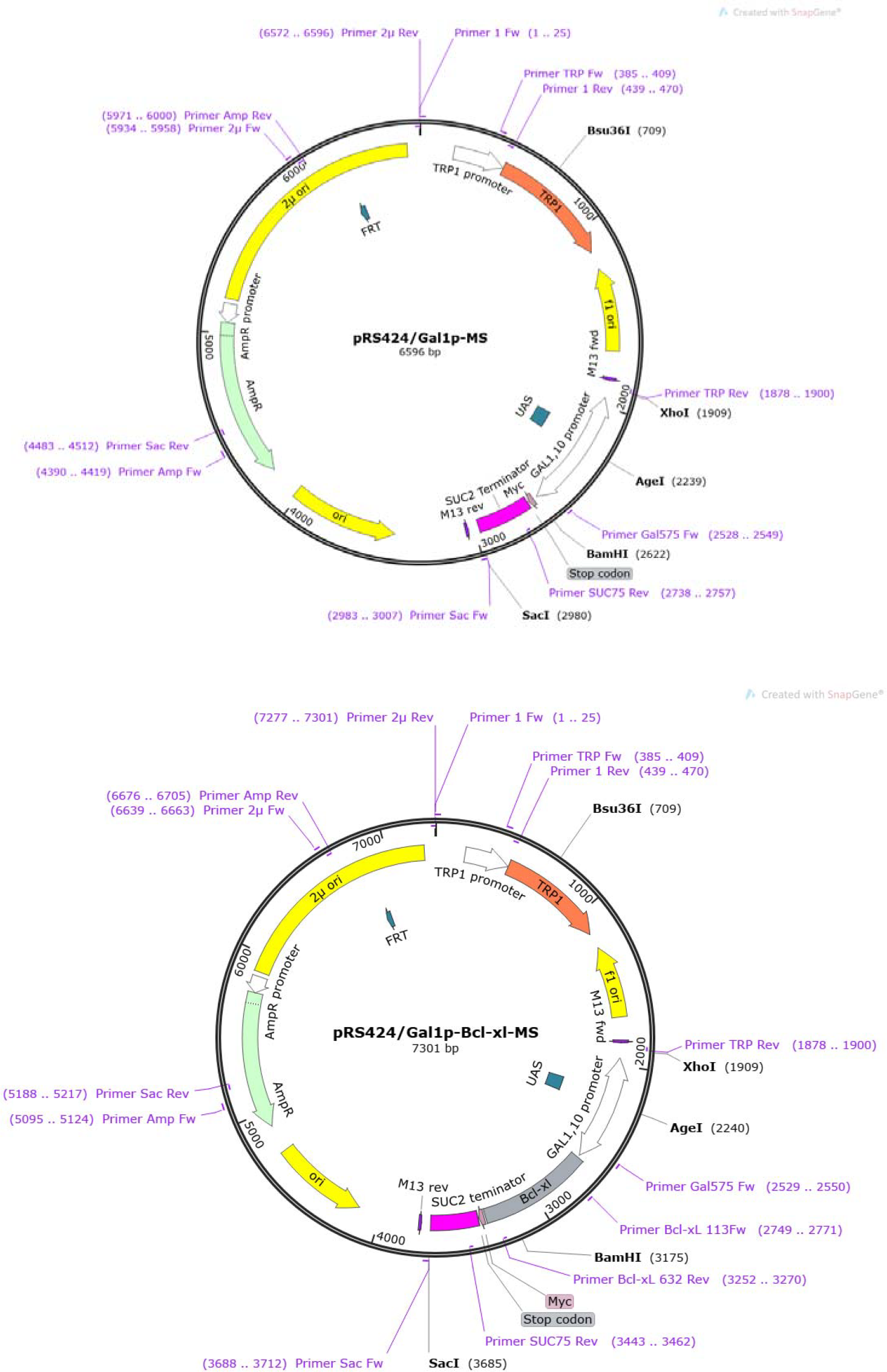
The location of primers for sequencing on new constructed pRS424

**Figure 16.**
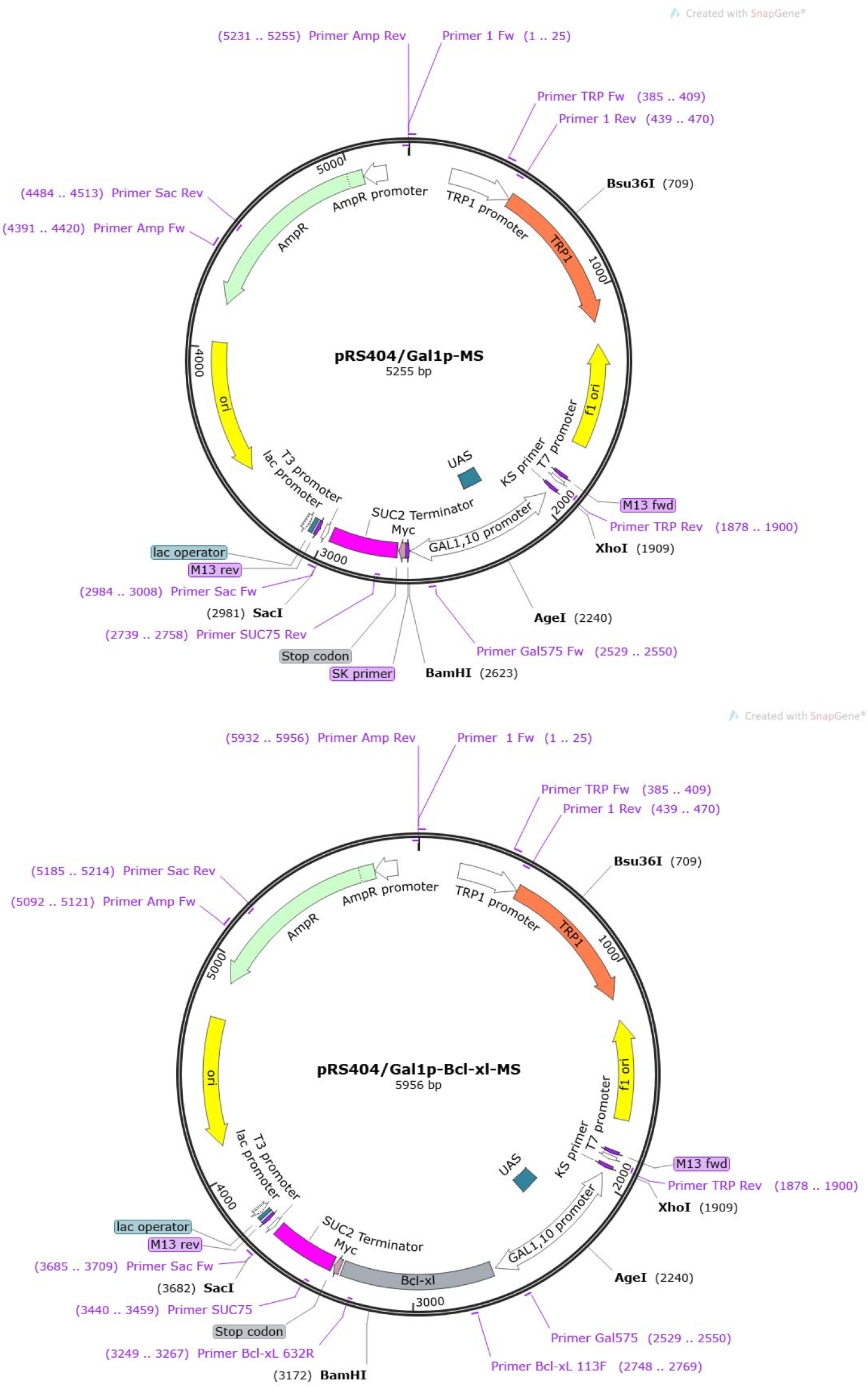
The location of primers for sequencing on new constructed pRS404.

**Table 2.**
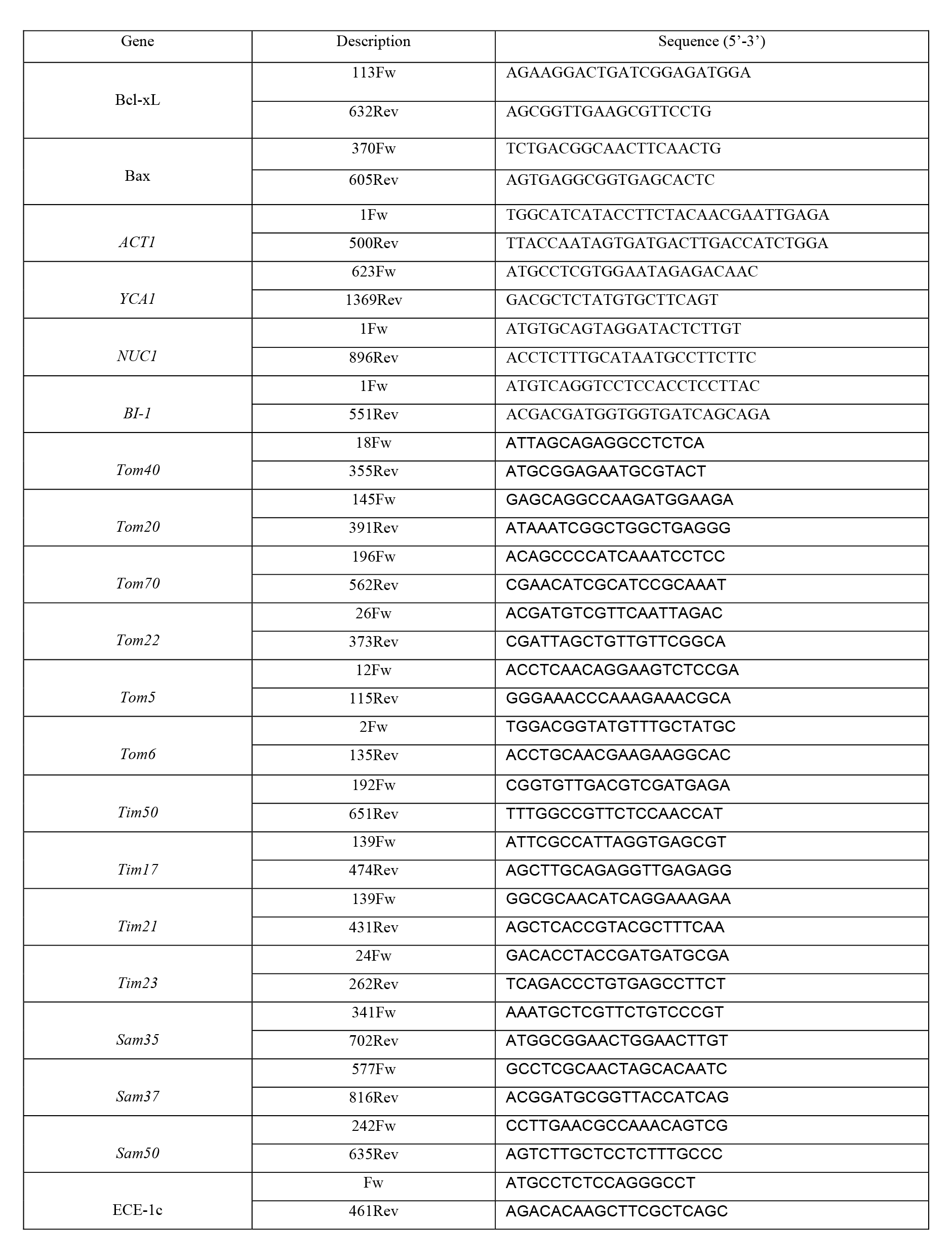
Primers used for detecting the expression level of candidate genes

The four new constructs were transformed into W303a and W303α strain respectively, the newly made strains (W303α: pRS404/Gal1p-MS, W303α: pRS404/Gal1p-Bcl-xL-MS, W303α: pRS424/Gal1p-MS, W303α: pRS424/Gal1p-Bcl-xL-MS) were cultured on one side of YPD agar plates. The strains after ECE-1 transformation, which contain both Bax and ECE-1 isoforms were cultured on the other side of YPD plates. Then the plates were incubated at 30°C overnight. The second day, to scratch cells from both side respectively, mix them in the middle of the plates to make two strains crossing at at 30°C overnight. The third day, to pick the middle cell of the plates then streak them onto SD-LEU-URA-TRP drop-out plates and incubated at 30°C for screening the single colonies of diploid cells. After 2 days, transfer the single colonies to new SD-LEU-URA-TRP drop-out plates and let them grow 1-2 days for obtaining the diploid strains, which contain Bax, ECE-1 isoforms and Bcl-xL in one cell.

### Colony PCR

Colony PCR was used for confirming the genotypes of new constructed strains. To make fresh solution 1B, pick a colony from plate and suspend it in 50 µl solution 1B, then 5 µl 10mg/ml Zymolyase was added. The mixture was incubated at 37°C for 30min and centrifuge at 8000rpm for 1min, then to discard the supernatant. Resuspend cell pellet with 50 µl 0.02M NaOH and heated at 100°C for 1min, the mixture can be used as the template for PCR.

### Spot assay

Spot assay was used to test the interaction between human Bax and candidate genes and observe the phenotype through the growth of yeast cells. The haploid cells or diploid cells contain human Bax, Bcl-xL and ECE-1 isoforms were screened by LEU-TRP, LEU-URA or LEU-URA-TRP drop-out media. The yeast cells were picked up into 500µl of sterilised distilled water and counted by haemocytometer to the same number of cells (10^6^ cells) in the first dilute of each sample. Then to make 10-fold serial dilutions (50 µl culture + 450 µl water) with 1.5ml micro tubes. The first group of tubes were undiluted, the second group is 10^-1^ dilution until to 10^-4^ dilution. 5 µl from each tube is spotted on SD-drop out as well SG-drop out plates. The spotted plates were left to dry for 10 minutes until the spots were absorbed thoroughly by the media, then incubated in 30°C for 48h.

### Growth curves recording of yeast strains

The ROTOR HDA system was purchased from Singer Instruments (Somerset, U.K) and the microplate reader was from BMG Labtech’s FLUOstar Omega, which is used for recording growth curves of yeast. A stock plate of the strains from yeast mating needs to be prepared. The cells of each strain were pelleted in 100 µl of YPD on a 96 well plate. The purpose of using YPD in this step is to let yeast cells grow faster and collect more cells than in the drop-out liquid media, though it might have a risk of plasmid loss. Then mix well and transfer them to a SD-URA-LEU-TRP drop-out plate by ROTOR. After incubated at 30 °C overnight, the strains were transferred to a soft YPD plate by the ROTOR for another day in 30 °C and keep the original plate maintained in 4°C as a stock plate. The final plate was spread with a SG-URA-LEU-TRP drop-out agar plate and incubated in microplate reader in 30°C for measuring the OD_600_ value and recording growth curves.

### Yeast DNA/RNA extraction, RT-PCR and Semi-qPCR

The yeast cells of each kind of strains were cultured and grown to exponential phase in liquid media with adding galactose (SG-LEU-URA-TRP), the cell pellet was harvest by centrifuging and can be stored directly at −80°C without adding glycerol, which is available for both DNA and RNA extraction.Total DNA of yeast strains was extracted by the classic phenol chloroform method with using Zymolyase for breaking yeast cell wall before the extraction. The concentration and value of A260/A280 of the DNA samples were measured by Nano Photometer, make aliquots of the rest of DNA samples (50 µl) and stored at −80°C for further experiments. Total RNA was extracted by using TRIzol Reagent (Thermofisher) based on the instruction. The RNA samples were purified by the DNase I (New England Biolabs) treatment according to the instruction after measuring concentration (µg/mL) and A260/A280. Finally, to make aliquots of 10 μl and measure the concentration and A260/A280 (the value between 1.8 and 2.0 shows a good quality of RNA samples). The purified RNA samples can be stored at −80°C, which were available for RT-PCR.

RT-PCR was used for confirming the expression of Bax-integrated yeast system in mRNA level. The cDNA samples were obtained from RT-PCR of purified RNA samples, which were purified by the gel recovery kit and diluted with 12 μl of elution buffer. To measure the concentration of purified cDNA samples by nanophotometer. Purified cDNA samples were stored at −80°C for the downstream semi-qPCR analysis. The semi-qPCR is a method for assessing the mRNA expression level of the candidate genes by measuring the grey value of the bands of PCR products. Gery value means the brightness of each band of a pixel, which is able to be measured by ImageJ. The candidate genes were set into 4 groups according to their function and location. The group 1 includes three yeast apoptotic factors, which includes YCA1 and NUC1 (pro-apoptotic genes) and BI-1 (anti-apoptotic gene) The group 2 includes the members of TOM complexes (two primary receptors of Tom70 and Tom22, the channel-forming protein Tom40, a secondary receptor of Tom22, two small subunits of Tom5 and Tom6). The members of SAM complexes (Sam50, Sam35, Sam37) were involved in the group 3. The translocases of inner membrane (two receptors of Tim50 and Tim21, two channel-forming proteins of Tim23 and Tim17) were included in the group 4.

### Data analysis

The plasmid maps of new constructs were generated from SnapGene Viewer 2.8.3, which was used for designing primers as well. All the data obtained from spot assay and semi-qPCR was analysed by ImageJ and Graphpad Prism 7.0. and presented as mean±SD, unless it was stated specifically. The statistical analysis was performed by One-way or Two-way analysis of variance (ANOVA) with Bonferroni’s post-test and the statistical significance was shown as *p<0.05, **p<0.01 or ***p<0.001.

### Data Availability

No data is deposited in external repositories in this study.

## Acknowledgement

We are much grateful to the research paper of Jing Lu and Gary Willars for the information of original vector plasmids for ECE-1 isoforms in cloning work. We thank Yishen Li for the help of operating ROTOR HDA colony manipulation robot for yeast work and Danae Georghiou for the assistance with PHENOS software. Thanks all the members in Centre of Genetic Architecture who have ever helped with the progress of the study.

## Author contributions

Conceptualization: All the authors; Investigation and methodology: Hanhui Xie and Yan Huang; Supervision: Edward J. Louis and Xiaodong Xie; Formal analysis and data curation: Hanhui Xie; Writing-original draft: Hanhui Xie; Writing-review and editing: All the authors.

